# Preventing trogocytosis by cathepsin B inhibition augments CAR T cell function

**DOI:** 10.1101/2024.06.11.598379

**Authors:** Kenneth A. Dietze, Kiet Nguyen, Aashli Pathni, Frank Fazekas, Wenxiang Sun, Ethan Rosati, Jillian M. Baker, Maday Galeana Figueroa, Etse Gebru, Daniel Yamoah, Rediet Mulatu, Alexander Wang, Aaron P. Rapoport, David Lum, Xiaoxuan Fan, Sabarinath V. Radhakrishnan, Djordje Atanackovic, Arpita Upadhyaya, Tim Luetkens

## Abstract

Chimeric antigen receptor (CAR) T cell therapy has shown remarkable efficacy in cancer treatment. Still, most patients receiving CAR T cells relapse within 5 years of treatment. CAR-mediated trogocytosis (CMT) is a potential tumor escape mechanism in which cell surface proteins transfer from tumor cells to CAR T cells. CMT results in the emergence of antigen-negative tumor cells, which can evade future CAR detection, and antigen-positive CAR T cells, which has been suggested to cause CAR T cell fratricide and exhaustion. Whether CMT indeed causes CAR T cell dysfunction and the molecular mechanisms conferring CMT remain unknown. Using a selective degrader of trogocytosed antigen in CAR T cells, we show that the presence of trogocytosed antigen on the CAR T cell surface directly causes CAR T cell fratricide and exhaustion. By performing a small molecule screening using a custom high throughput CMT-screening assay, we found that the cysteine protease cathepsin B is essential for CMT and that inhibition of cathepsin B is sufficient to prevent CAR T cell fratricide and exhaustion, leading to improved long-term CAR T cell persistence and anti-tumor activity. Our data demonstrate that it is feasible to separate CMT from cytotoxic activity, that CAR T cell persistence, a key factor associated with clinical CAR T cell efficacy, is directly linked to cathepsin B activity in CAR T cells, and that it is possible to improve CAR T cell function through selective inhibition of CMT.

<u>One sentence summary:</u> CAR-mediated trogocytosis is mediated by the cysteine protease cathepsin B and directly causes CAR T cell exhaustion and fratricide.

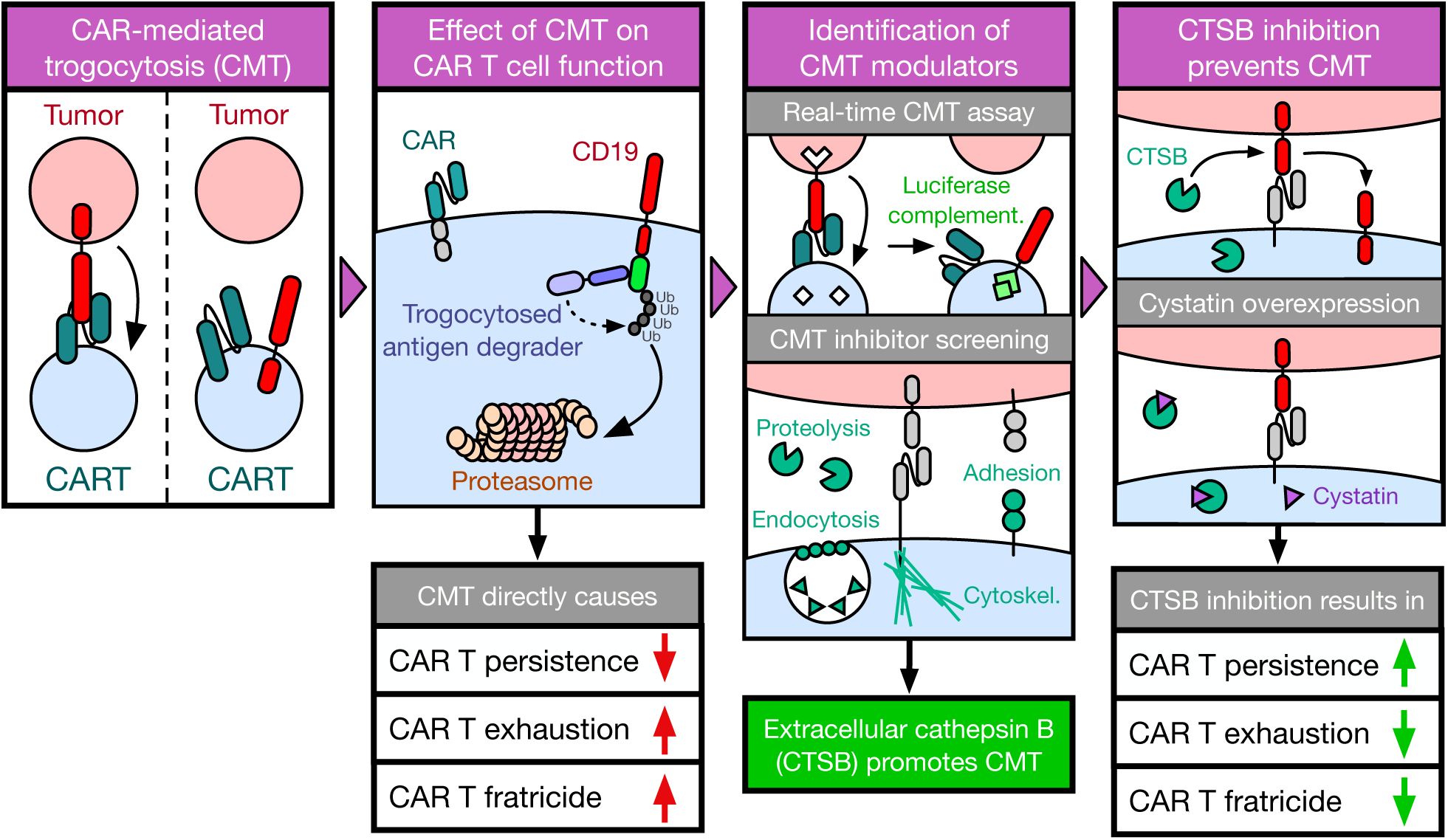

## INTRODUCTION

Chimeric antigen receptors (CAR) are genetically engineered proteins combining antigen binding domains with immune cell activation domains^1–3^. T cells and natural killer (NK) cells can be engineered to express CARs to recognize tumor-associated antigens with high specificity and have been shown to be clinically effective at targeting several hematological malignancies^1,4–7^. Currently, seven CAR T cell approaches targeting CD19 or B cell maturation antigen (BCMA) have been approved in the US for the treatment of hematological cancers^8,9^. While CAR T cell therapy has greatly improved clinical outcomes for patients, many patients still relapse within five years of treatment^10–15^. Several factors have been implicated in CAR T cell-associated relapse, including increased CAR T cell exhaustion^16^, reduced CAR T cell expansion^17^ and persistence^18,19^, as well as tumor antigen escape^18,2018,21^.

It has previously been shown that CAR T cells and CAR NK cells rapidly transfer the targeted antigen from tumor cells to their own cell surface in a process called trogocytosis^22–24^. Trogocytosis was initially observed at the immune synapse between conventional T cells and antigen presenting cells (APCs), where TCR internalization following activation resulted in the transfer of MHC to recipient T cells^25,26^. CAR-mediated trogocytosis (CMT) occurs in both hematological and solid tumors and is correlated with increased expression of apoptotic and exhaustion markers on CAR T cells, as well as reduced persistence and expansion^22–24^. In addition, it has been shown that T cells engineered to express the tumor antigen CD19 are efficiently killed by other CD19 CAR T cells (“fratricide”)^22^. CMT is also implicated in a reduction of target antigen on the tumor cell surface, leading to antigen-negative or antigen-low variants which could escape CAR T cell detection^23^. Reductions in CAR T cell affinity have previously been shown to reduce CMT^23^, but this could lead to an increased antigen threshold^27^, preventing the detection of antigen-low tumor cells.

Mechanistically linking CMT to exhaustion, fratricide, and reduced CAR T cell persistence has, so far, been elusive due to the lack of specific inhibitors of trogocytosis. In this study, we develop an analytical framework to investigate CMT and determine the effects of CMT on CAR T cell function as well as its mechanistic basis. In addition, we demonstrate how CMT can be specifically targeted by inhibiting the protease cathepsin B using endogenous inhibitors to increase CAR T cell persistence and enhance long-term anti-tumor activity.

## RESULTS

### CAR T cells acquire target antigen via trogocytosis

CAR-mediated trogocytosis (CMT) is the extraction of the targeted tumor antigen from the tumor cell surface and its incorporation into the CAR T cell plasma membrane^22–24,28^ (Suppl. Fig. S1A). *Ex vivo*, CMT occurs across cancer types^22,23^ and target antigens, including CD19^22–24^, CD22^22^, mesothelin^22^, and BCMA^28^. We found robust transfer of CD19 to CD19-targeting CAR T cells (clone: FMC63, Suppl. Fig. S1B) as well as CD19 loss on tumor cells (Suppl. Fig. S1C), which can be observed minutes after CAR T cells contact target cells (Fig. 1A, Suppl Fig. S1D). Similarly, we confirmed transfer of *B cell maturation antigen* (BCMA), the target of two clinically approved CAR T cell approaches for the treatment of multiple myeloma^4,5,29,30^ (Fig. 1B) to BCMA CAR T cells (Fig. 1C) and loss of BCMA from tumor cells (Fig. 1D) independent of BCMA binding domain (Fig. 1B). When assessing CMT in solid tumor models *in vitro*, we observed significant transfer of target antigen to CAR T cells targeting LINGO1 (Suppl. Fig. S1E), GD2 (Suppl. Fig. S1F), and FolR⍺ (Suppl. Fig. S1G). In the clinical setting, CMT has only been shown to occur in B cell lymphoma patients treated with CAR NK cells^24^. By analyzing peripheral blood mononuclear cells (PBMCs) from patients who had recently received CD19 CAR T cells, we demonstrate that CMT also occurs in some patients treated with CAR-expressing T cells (Fig. 1E/F, Suppl. Fig. S1H). Further analyzing a sample from a patient with a high number of CD19^+^ CAR T cells, we found that the presence of CD19 on CAR T cells was associated with increased amounts of the exhaustion markers PD-1 and TIM-3 (Suppl. Fig. S1I).

**Figure 1:**
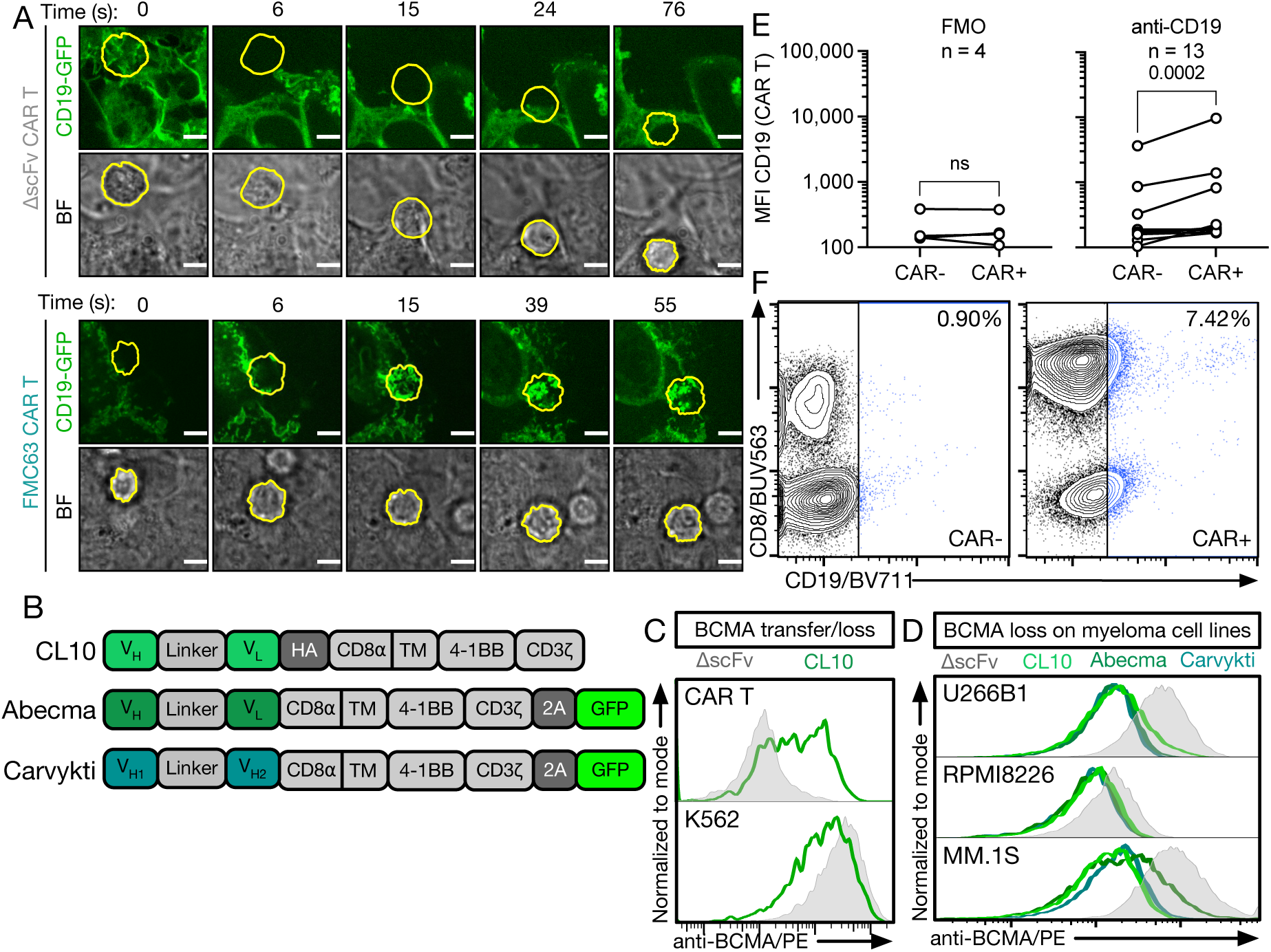
CD19 and BCMA CAR T cells rapidly acquire target antigen via trogocytosis. **(A)** Observation of CMT in brightfield (BF) and confocal images of ΔscFv or FMC63 CAR T cells. CAR T cells were cultured with plate-bound 293T cells expressing a CD19-GFP fusion protein. CD19 is shown in green. Individual CAR T cells are circled in yellow and tracked over time. Scale bar represents 5µm. Data is representative of cells from two independent experiments. **(B)** Schema of BCMA-targeting CAR T cell products. The same PBMC donor was used to produce Abecma and Carvykti CAR T cells; due to availability, a separate donor was used to produce CL10 CAR T cells. **(C)** BCMA transfer to CAR T cells (top) and BCMA loss on BCMA-GFP-expressing K562 cells (bottom) as determined by flow cytometry. **(D)** BCMA loss on MM.1S, RPMI8226, and U266B1 cells when cocultured with CL10^31^, Abecma, and Carvykti CAR T cells as determined by flow cytometry. **(C/D)** Data is representative of three independent experiments. **(E)** Amount of CD19 on CAR T cells (CAR^+^) or non-CAR T cells (CAR^-^) in PBMCs isolated from patients 7-28 days after receiving CD19 CAR T cell therapy using full-spectrum flow cytometry. Data indicate values from individual patient samples (FMO n = 4; anti-CD19 n = 13). Statistical significance was determined by Wilcoxon matched-pairs signed-rank test. Fluorescence-minus-one (FMO) refers to staining with full flow cytometry panel except anti-CD19. **(F)** Representative flow cytometry data from one patient depicting CD19 levels on CAR^-^ and CAR^+^ T cells. CAR was stained using an anti-idiotype antibody.

### CAR-mediated trogocytosis directly causes CAR T cell dysfunction

The presence of trogocytosed antigen on CAR T cells is correlated with increased exhaustion and reduced viability^22–24^ but it remains unknown if CMT directly causes this T cell dysfunction. To answer this question, we developed an approach for the targeted degradation of trogocytosed CD19 fused to GFP (CD19-GFP) in CAR T cells (Fig. 2A). To validate the approach, we first expressed CD19-GFP as well as a trogocytosed antigen degrader (TAD), the E3-targeting domain Nslmb^32,33^ fused to a GFP-binding protein (TAD_GFP_), in 293T cells. TAD overexpression resulted in a significant reduction of total CD19, with multiple CD19 bands likely reflecting differential glycosylation^34,35^ (Suppl Fig. S2A), and surface CD19-GFP (Suppl Fig. S2A/B). We next generated CAR T cells expressing TAD_GFP_ and did not observe an effect of TADon CAR T cell expansion or short-term antitumor activity (Suppl Fig. S2C/D).

**Figure 2:**
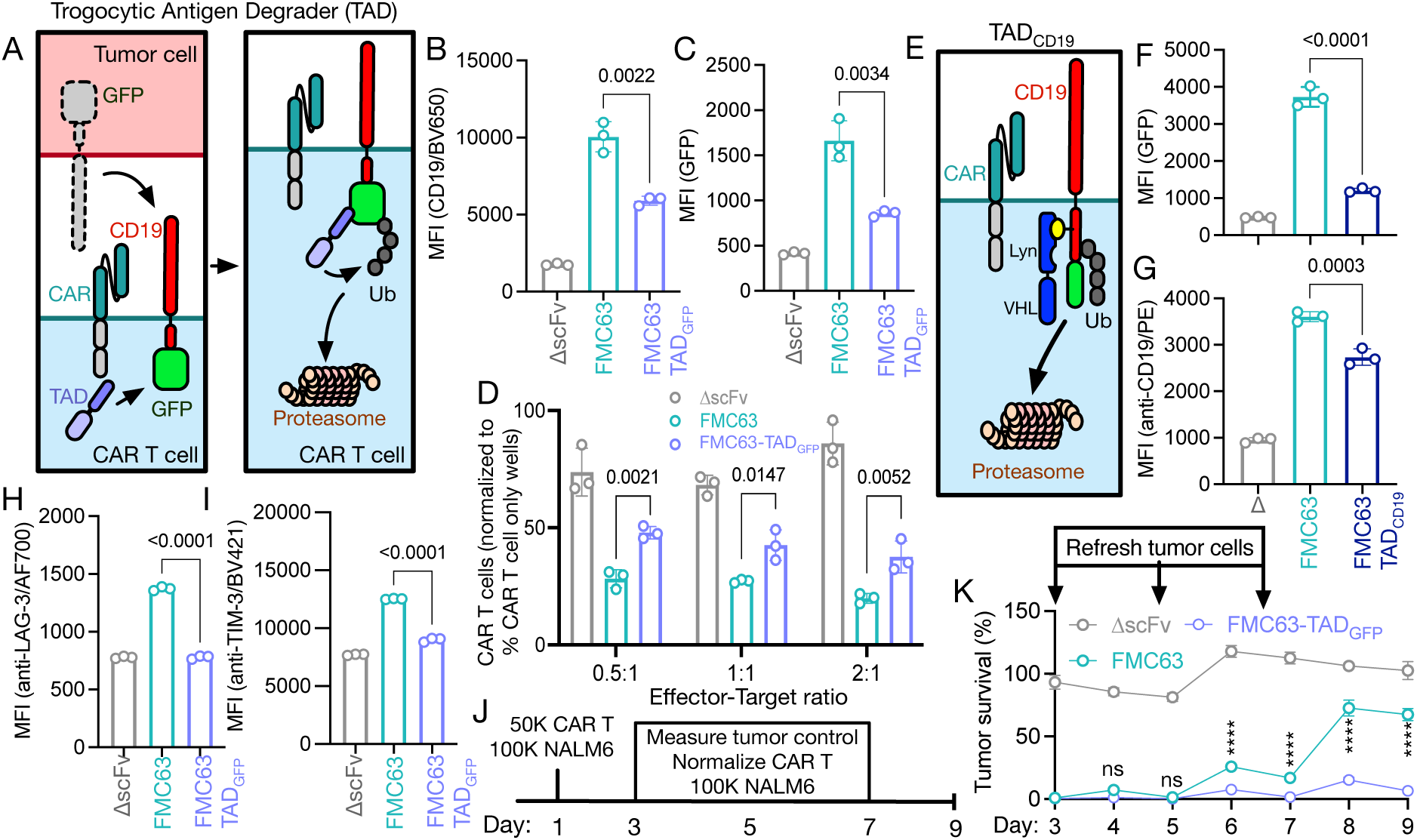
CAR-mediated trogocytosis directly causes CAR T cell dysfunction. **(A)** Schema of trogocytosed antigen degrader (TAD). **(B)** CD19 and **(C)** GFP expression on FMC63 CAR T cells expressing TADGFP or T cells expressing a CAR lacking a binding domain (ΔscFv) following a 2-hour coculture with CD19-GFP-expressing NALM6 cells as determined by flow cytometry. Data indicates mean ± S.D. from three technical replicates. **(D)** Quantification of CAR T cells after a 24-hour coculture with CD19-GFP-expressing NALM6 cells using flow cytometry. Values are normalized to wells containing only CAR T cells. **(B-D)** Data indicates mean ± S.D. from three technical replicates. Data is representative of three independent experiments. **(E)** Schema of a fully human-derived TAD system (TADCD19). Lyn = SH2 domain of lyn kinase; VHL = von Hippel-Lindau protein; Ub = ubiquitin. **(F)** GFP and **(G)** CD19 expression on FMC63 CAR T cells expressing TADCD19 or ΔscFv CAR T cells after a 2-hour coculture with CD19-GFP-expressing NALM6 cells at a 0.5:1 effector-target ratio. Data indicates mean ± S.D. from three technical replicates. Data is representative of two independent experiments. **(H)** LAG-3 and **(I)** TIM-3 expression on CAR T cells expressing TADGFP as measured by MFI after a 24-hour coculture with CD19-GFP-expressing NALM6 cells using full-spectrum flow cytometry. Data indicates mean ± S.D. from three technical replicates. Data is representative of three independent experiments. **(B-D, F-I)** Statistical significance was determined by one-way ANOVA. **(J)** Schema of *in vitro* serial coculture experiment to determine long-term CAR T cell expansion and antitumor activity. CAR T cells were normalized and fresh tumor cells were added on days 3, 5, and 7. **(K)** Survival of CD19-GFP-expressing NALM6 cells expressing firefly luciferase (Fluc) at a 0.5:1 effector-target ratio using a luminescence-based cytotoxicity assay. Data indicates mean ± S.D. from three technical replicates. Tumor survival was normalized to untreated tumor cells. Data is representative of two independent experiments. Statistical significance comparing FMC63 and FMC63-TADGFP was determined by two-way ANOVA.

When cocultured for 2 hours with K562 cells expressing CD19-GFP, conventional FMC63 CAR T cells showed increased levels of CD19 (Fig. 2B, Suppl. Fig. S2E) and GFP (Fig. 2C, Suppl. Fig. S2F) indicative of CMT, as expected. However, FMC63 CAR T cells also expressing TAD_GFP_ showed an approximately 50% reduction in both CD19 and GFP after coculture. Using the TAD system, we next investigated the effect of CMT on CAR T cell fratricide. We found that TAD_GFP_ expression did not alter the proliferation of CAR T cells when co-cultured with CD19^+^ tumor cells (Suppl. Fig. S2G). However, TAD_GFP_ expression resulted in a two-fold increase in CAR T cell numbers after co-culture (Fig. 2D), indicating that CMT directly causes CAR T cell death, likely due to fratricide. Targeted degradation of trogocytosed antigen was also validated using a fully human TAD (TAD_CD19_) construct directly targeting CD19. TAD_CD19_ recognizes CD19 by using the Lyn kinase SH2 domain (Fig. 2E), which has been shown to bind the intracellular domain of CD19 with high affinity^36^. Co-expression of TAD_CD19_ in FMC63 CAR T cells also resulted in reduced CMT (Fig. 2F/G). The reduction in CD19-GFP on the CAR T cell surface by TAD_CD19_ was more pronounced when measuring GFP (Fig. 2F) than CD19 (Fig. 2G), which may be the result of CD19 CAR molecules binding to the trogocytosed CD19 in cis blocking the anti-CD19 detection antibody and thereby leading to an underestimation of CD19 on the CAR T cell surface. These data indicate that selective degradation of trogocytosed antigen is feasible in CAR T cells and that this approach can be used to explore the effects of CMT on CAR T cell function.

It has previously been shown that FMC63 binding to CD19 in *cis*^37^ can cause persistent CAR signaling followed by T cell exhaustion. We found that conventional FMC63 CAR T cells that had acquired CD19 during co-culture with CD19-GFP-expressing K562 cells showed a two-fold increased expression of exhaustion markers LAG-3 and TIM-3 compared to negative control T cells expressing a CAR without a binding domain (ΔscFv, Fig. 2H/I, Suppl. Fig. S2H/I). Expression of TAD_GFP_ prevented LAG-3/TIM-3 upregulation on these cells, demonstrating that increased exhaustion marker expression is the direct consequence of trogocytic antigen transfer to CAR T cells. CMT-induced fratricide and exhaustion could substantially limit long-term CAR T cell anti-tumor activity. In a serial coculture stress test of CAR T cell efficacy (Fig. 2J), we found that TAD_GFP_ substantially increased the ability of CAR T cells to control B cell acute lymphoblastic leukemia (B-ALL, Fig. 2K).

These data demonstrate that CMT directly causes CAR T cell dysfunction and fratricide, thereby limiting CAR T cell persistence and anti-tumor activity.

### Real-time detection of CAR-mediated trogocytosis by luciferase complementation assay

Little is known regarding the molecular and cellular drivers of CMT, and to date there are no high-throughput assays to systematically probe potential modulators of CMT as it occurs in real-time. To address this, we developed a luciferase complementation assay for the real-time detection of CMT (CompLuc, Fig. 3A). In the CompLuc assay, luciferase complementation occurs following the transfer of CD19 fused to the C-terminal fragment of NanoLuc^38^ (cLuc) to CAR T cells, which express the complementary cytosolic N-terminal NanoLuc (nLuc) fragment with high affinity for cLuc^38^ (Fig. 3B, Suppl. Fig. S3A/B). The CompLuc assay allows for sensitive detection of small numbers of nLuc^+^cLuc^+^ CAR T cells in cocultures (Fig. 3C). We demonstrate that CMT, as quantified by CompLuc, is antigen-and CAR-dependent, and correlates with effector-target ratio (Fig. 3D-F) and antigen transfer as measured by flow cytometry (Fig. 3D/F).

**Figure 3:**
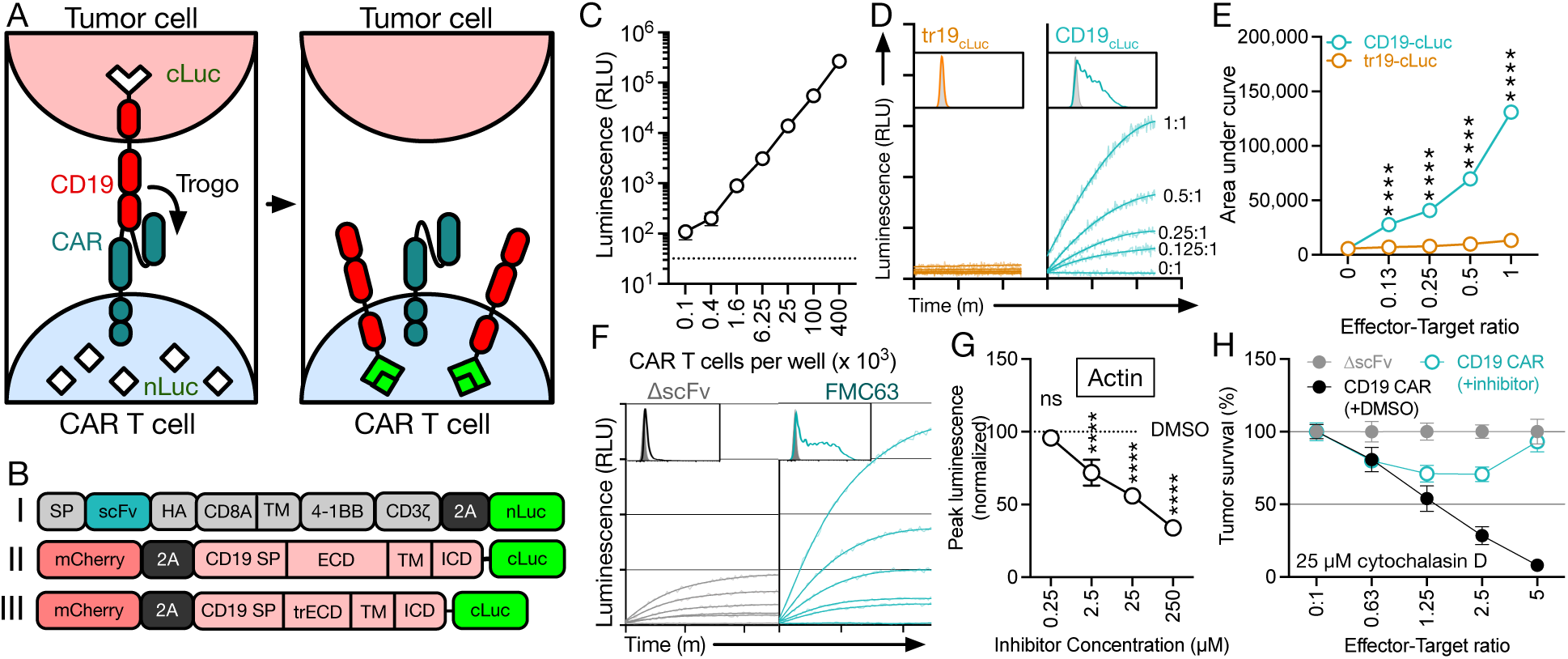
Luciferase complementation assay allows for the real-time detection of trogocytosis. **(A)** Schema of luciferase complementation assay (CompLuc). **(B)** Schema of constructs used for CAR T cells (I) and tumor cells (II/III) for development of luciferase complementation assay. SP = signal peptide; TM = transmembrane region; trECD = truncated extracellular domain; ECD = extracellular domain; ICD = intracellular domain. **(C)** Luminescence of nLuc+cLuc+ CAR T cells. Data represent mean ± S.D. of three technical replicates and is representative of two independent experiments. **(D)** CMT during coculture of FMC63 CAR T cells expressing nLuc with K562 cells expressing full length (teal) or truncated (tr19, orange) CD19 fused to cLuc using CompLuc or flow cytometry after 1 hour at the indicated effector-target ratios. CMT using CompLuc was measured for a total of 2.5 hours at 1-minute intervals. Data represents best fit of three technical replicates. Boxed area represents histogram of anti-CD19/FITC staining of CAR T cells at a 1:1 effector-target ratio after 1 hour as measured by flow cytometry. **(E)** Area-under-curve quantification of luminescence in Fig. 2D. Data represent mean ± S.D. of three technical replicates. **(F)** CMT during coculture of ΔscFv or FMC63 CAR T cells expressing nLuc with K562 cells expressing full length CD19-cLuc using CompLuc or flow cytometry after 1 hour at the indicated effector-target ratios. CMT using CompLuc was measured for a total of 2.5 hours at 1-minute intervals. Data represents best fit of three technical replicates. Boxed area represents histogram of anti-CD19/FITC staining of CAR T cells at a 1:1 effector-target ratio after 1 hour as measured by flow cytometry. **(D-F)** Data is representative of at least three independent experiments. **(G)** Peak luminescence of FMC63 CAR T cells cocultured with K562 cells expressing CD19-cLuc at a 1:1 effector-target ratio as determined by CompLuc. CAR T cells were pre-treated with cytochalasin D for 1 hour at the indicated concentrations. **(H)** Survival of Raji-Fluc cells after a 16-hour coculture with FMC63 CAR T cells pre-treated with the indicated concentrations of cytochalasin D. CMT values are normalized to a DMSO control. **(G/H)** Data represent mean ± S.D. of three technical replicates. Data is representative of two independent experiments.

To validate the CompLuc assay, we determined the effect of an established modulator of CMT, the actin polymerization inhibitor cytochalasin D^22,39^. Treatment of CAR T cells with cytochalasin D significantly reduced CMT in a dose-dependent manner as measured by CompLuc (Fig. 3G), demonstrating the ability of the assay to identify modulators of CMT.

### Extracellular cathepsin B is a key driver of CAR-mediated trogocytosis

CMT may be the result of incomplete target cell killing followed by the endocytosis of target cell fragments, as suggested by the simultaneous effect of cytochalasin D on CMT and CAR-mediated cytotoxicity (Fig. 3H). However, we hypothesized that, while the molecular mechanisms driving target cell killing and trogocytosis may overlap, CMT could in part be mechanistically distinct from cytotoxic function.

To identify functional systems that could be specifically involved in CMT but not cytotoxicity, we explored the effect of inhibiting various processes at the T cell immune synapse on CMT^22,39–42^ (Fig. 4A). We found that inhibition of cathepsin B (CTSB) with the small molecule inhibitor Ca-074-Me in FMC63 CAR T cells reduced CMT in a dose-dependent manner as measured by CompLuc (Fig. 4B) without substantially altering CAR T cell cytotoxicity (Fig. 4C). Using a membrane-impermeable inhibitor of CTSB (Ca-074), we found that inhibition of extracellular CTSB alone is sufficient to prevent CMT (Fig. 4D).

**Figure 4:**
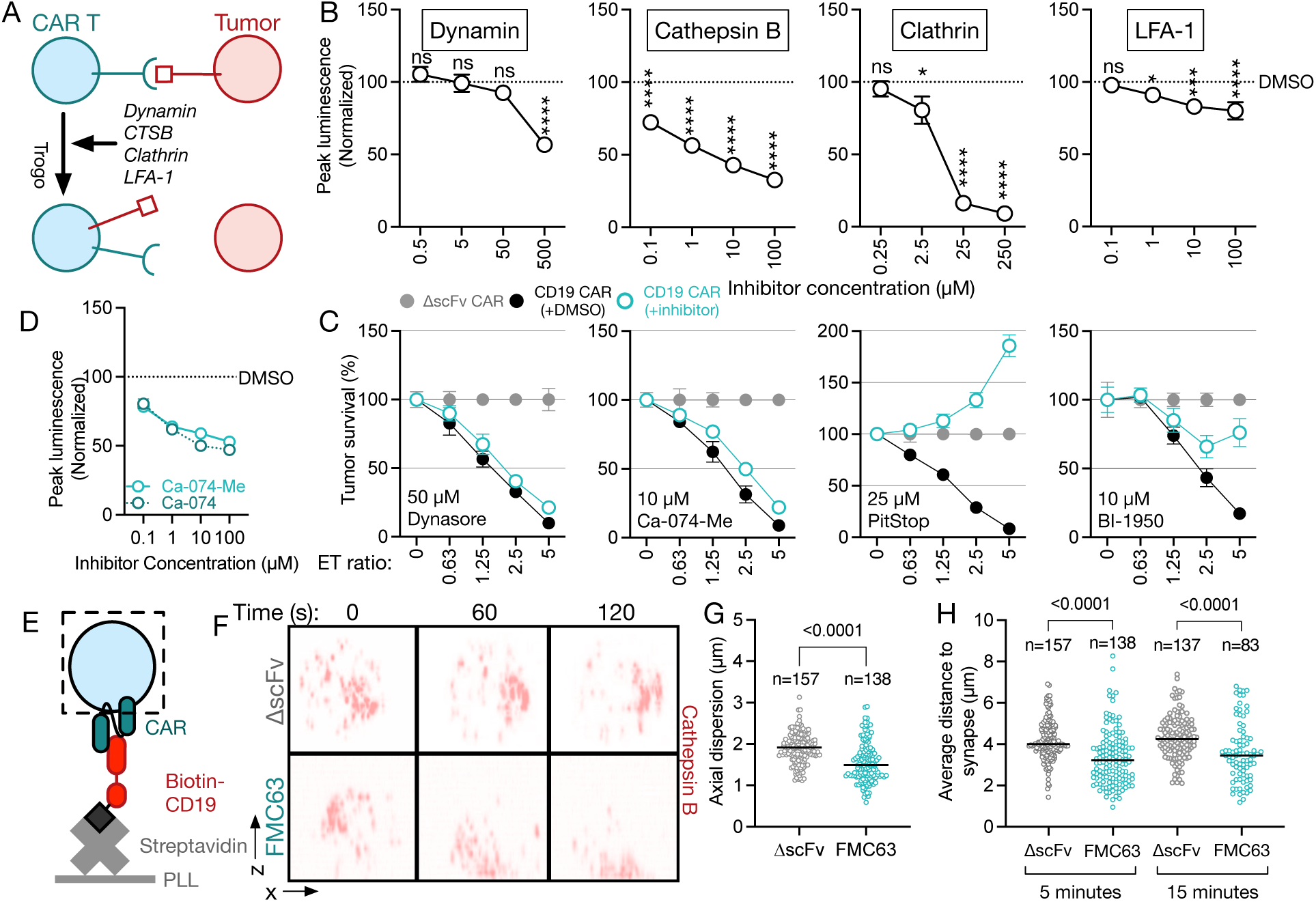
Extracellular cathepsin B is a key driver of CAR-mediated trogocytosis. **(A)** Schema of potential modulators of CMT. **(B)** Peak luminescence as determined by CompLuc of FMC63 CAR T cells treated with inhibitors of the indicated proteins cocultured with K562 cells expressing CD19-cLuc at a 1:1 effector-target ratio. Values are normalized to DMSO/vehicle control. Data represent mean ± S.D. of three technical replicates. Data is representative of three independent experiments. **(C)** Survival of Raji-Fluc cells after 16-hour coculture with FMC63 CAR T cells treated with inhibitors of the indicated proteins at a single concentration. Tumor survival was measured using a luciferase-based cytotoxicity assay. Data represent mean ± S.D. of three technical replicates. Data is representative of two independent experiments. **(D)** Peak luminescence as determined by CompLuc of FMC63 CAR T cells cocultured with K562 cells expressing CD19-cLuc at a 1:1 effector-target ratio. FMC63 CAR T cells were pre-treated with the indicated concentrations of Ca-074-Me (membrane-permeable) or Ca-074 (membrane-impermeable). Values are normalized to DMSO/vehicle control. Data represent mean ± S.D. of three technical replicates. Data is representative of three independent experiments. **(E)** Schema of imaging assay setup used to assess cathepsin B (CTSB) localization to the immune synapse. Biotinylated CD19 was immobilized on plate-bound NeutrAvidin. ΔscFv or FMC63 CAR T cells expressing CTSB-mCherry were added to slides and imaged using confocal microscopy. **(F-H)** ΔscFv or FMC63 CAR T cells expressing CTSB-mCherry were imaged by spinning disk confocal imaging after exposure to CD19 immobilized on glass slides at different time points. **(F)** Side (x-z) view of the distribution of CTSB-mCherry (red) at the indicated time points. The bottom of each box is aligned to the interface between the glass slide and the T cell. **(G)** Axial dispersion or **(H)** average distance of CTSB to the site of antigen contact. Data represent values from individual cells fixed 5 or 15 minutes after exposure to recombinant CD19. Numbers represent cells analyzed for each condition. **(G/H)** Statistical significance was determined by two-tailed Student’s *t*-test. **(F-H)** Data is representative of at least two independent experiments.

CTSB is a ubiquitously expressed cysteine protease, primarily localized to lysosomal and endosomal compartments under physiological conditions, where it is primarily involved in protein degradation and turnover^43,44^. However, CTSB is also found in the cytosol and exocytic vesicles, retains its catalytic activity at neutral pH^45^, and has been shown to line exocytic granules of cytotoxic T lymphocytes^46^. In addition to proteolytic degradation within the endosome and lysosome, CTSB has been implicated in the degradation and remodeling of components of the extracellular matrix, such as collagen and fibronectin, thereby promoting the invasion and metastasis of tumor cells^45,47,48^. Similarly, CTSB may contribute to the extraction of antigen-rich membrane fragments of target cells prior to their transfer to CAR T cells.

To further support the potential role of extracellular CTSB produced by CAR T cells in CMT, we next determined whether CTSB in CAR T cells is actively transported to the immune synapse upon antigen contact. To this end, we generated FMC63 CAR T cells expressing CTSB fused to mCherry and exposed these cells to immobilized CD19 (Fig. 4E). Specifically, we assessed intracellular CTSB trafficking in these cells using live-cell confocal imaging after exposing CAR T cells to recombinant CD19 immobilized on a glass slide. When assessing CTSB movement, we found that CTSB in FMC63 CAR T cells, but not in ΔscFv CAR T cells, rapidly localized to the site of antigen contact (Fig. 4F, Suppl. Fig. S1D), correlating with the timing of CMT as measured by CompLuc and flow cytometry (Fig. 3F). Specifically, following exposure to CD19, CTSB in FMC63 CAR T cells was significantly less dispersed throughout the cell (“axial dispersion,” Fig. 4G) indicating coordinated CTSB trafficking and localization significantly closer to the site of antigen contact (Fig. 4H) compared to ΔscFv CAR T cells. These findings suggest that CTSB is actively and rapidly transported to the immune synapse upon antigen contact and that its subsequent secretion by CAR T cells contributes to CMT.

### Cystatin abundance controls CTSB activity and CMT

CTSB is ubiquitously expressed across tissues and cell types, and its activity is regulated through a network of endogenous protease inhibitors called cystatins^49^. Cystatin A (CSTA) has been described as the main regulator of CTSB activity^50^. Like Ca-074, CSTA inhibits CTSB through insertion of a hydrophobic wedge into the active-site cleft of CTSB^51^ (Fig. 5A). To determine if CSTA abundance directly regulates CMT in CAR T cells, we engineered FMC63 CAR T cells to stably overexpress CSTA (CAR_CSTA_, Fig. 5B). We confirmed that CSTA overexpression led to significantly increased levels of CSTA protein in CAR T cells (Fig. 5C), which resulted in reduced CTSB activity (Fig. 5D) without altering *in vitro* CAR T cell expansion (Suppl. Fig. S4A/B), phenotype (Suppl. Fig S4C/D), or cytokine secretion (Suppl. Fig. S4E). It has previously been shown that CTSB^KO^ mice are viable, suggesting that loss of CTSB activity may not cause major toxicity^52–55^ . To determine if CSTA overexpression by engineered T cells causes toxicity, we treated NRG mice with CAR T cells overexpressing human CSTA (hCSTA) or mouse CSTA (mCSTA, Suppl. Fig. S5A-D) and found that CSTA overexpression did not result in weight loss or changes in major tissue architecture (Suppl. Fig. S5E-F). When co-cultured with tumor cells, CD19 CAR_CSTA_ cells showed equivalent short-term anti-tumor activity compared to conventional CAR T cells (Fig. 5E, Suppl. Fig. S6A) whereas BCMA CAR_CSTA_ cells showed slightly improved short-term anti-tumor activity (Suppl. Fig. S6B). When co-cultured with CD19^+^ or BCMA^+^ K562 cells, we observed significantly reduced CMT in CAR_CSTA_ cells as measured by CompLuc (Fig. 5F/G, Suppl. Fig. S6C/D) despite comparable CAR T cell activation (Suppl. Fig. S6E). CMT was similarly reduced when overexpressing a truncated CSTA_1-57_ containing the hydrophobic wedge (Suppl. Fig S6F) or the closely related protein cystatin B^56^ (Suppl. Fig S6G-I). CAR_CSTA_ T cells cocultured with relevant hematologic or solid tumor cell lines showed reduced antigen transfer to CAR T cells (Fig. 5H-J, Suppl. Fig. S6J-M) and antigen loss on tumor cells (Fig. 5K-M, Suppl. Fig. S6N), suggesting inhibition of CMT at the antigen extraction step. This CSTA-mediated inhibition of CMT resulted in significantly increased CD19 and BCMA CAR T cell numbers after short-term exposure to tumor cells *in vitro* (Fig. 5N/O, Suppl. Fig. S6O). To explore whether CSTA overexpression would be effective in a primary tumor cell model, we cocultured CAR or CAR_CSTA_ T cells with primary tumor cells from a patient with B-cell acute lymphoblastic leukemia (B-ALL) and found that CSTA overexpression significantly reduced CMT (Fig. 5P/Q) and increased CAR T cell numbers (Fig. 5R) in this setting as well. These data demonstrate that, in short-term co-cultures with tumor cells, CSTA overexpression significantly reduces CMT and increases CAR T cell survival.

**Figure 5:**
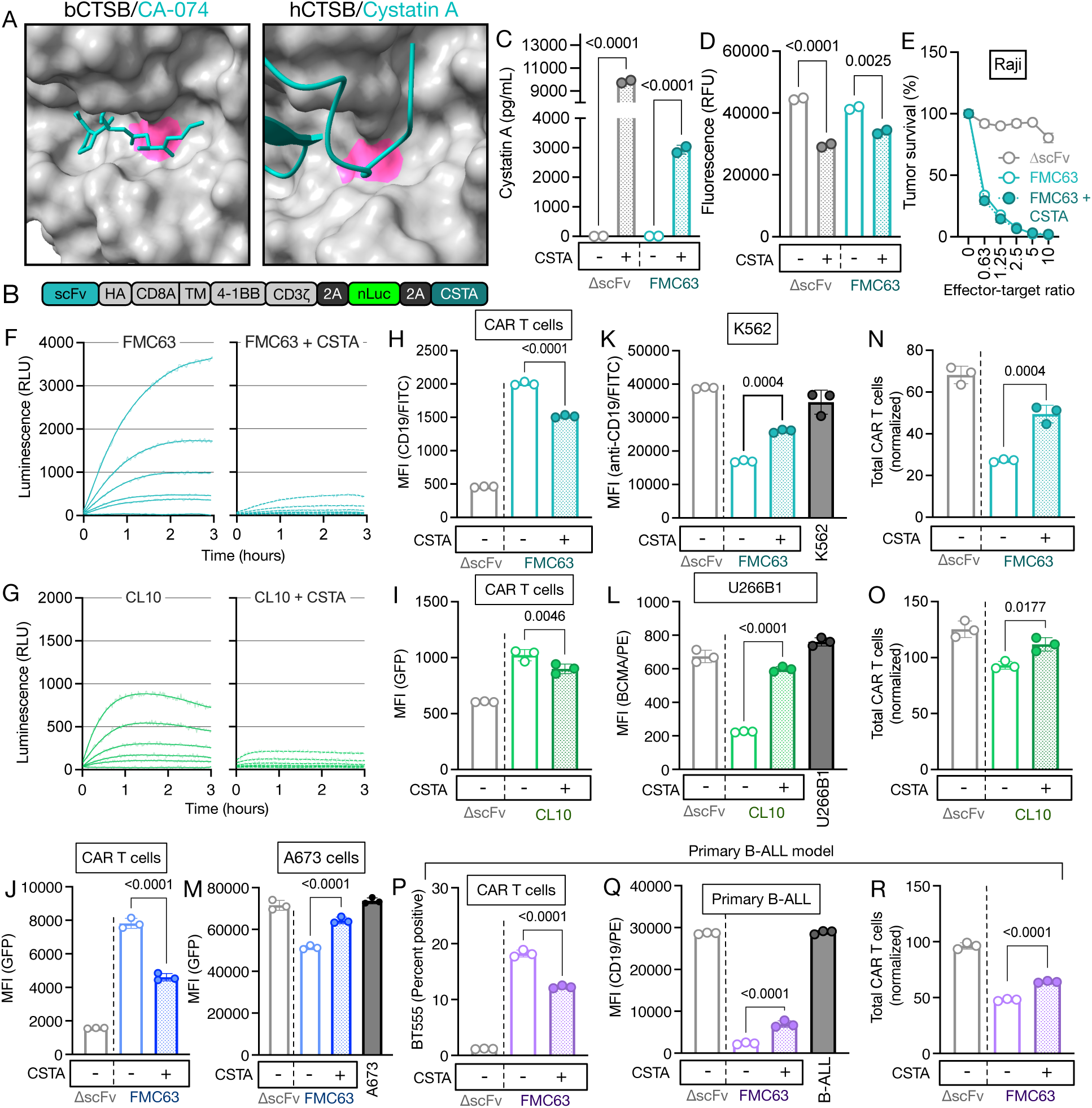
Cystatin abundance controls CTSB activity and CMT. **(A)** Crystal structures of bovine cathepsin B in complex with Ca-074 (left, PDB: 1QDQ) and human cathepsin B in complex with cystatin A (right, PDB: 3K9M). The active site cysteine of cathepsin B is shown in pink; Ca-074 and cystatin A are shown in teal. **(B)** Schema of construct used for production of CARCSTA T cells. **(C)** Levels of CSTA in CAR or CARCSTA T cells measured by ELISA. Data represent mean ± S.D. of two technical replicates. **(D)** Cathepsin B activity in CAR or CARCSTA T cells measured by fluorescence-based CTSB activity assay. Data represent mean ± S.D. of two technical replicates. **(C/D)** Data is representative of two independent experiments. **(E)** Survival of Raji-Fluc cells after a 16-hour coculture with FMC63 CAR or CARCSTA T cells at the indicated effector-target ratios. Data represent mean ± S.D. of three technical replicates. Data is representative of at least three independent experiments using cells produced from three healthy donors. **(F)** CMT of FMC63 CAR or CARCSTA T cells as determined by CompLuc using K562 cells expressing CD19-cLuc after 3 hours. Data is representative of three independent experiments using cells produced from three independent donors. **(G)** CMT in BCMA CAR or CARCSTA T cells as determined by CompLuc using K562 cells expressing BCMA-GFP-cLuc after 3 hours. Data is representative of three independent experiments. **(F/G)** Data represents best fit of three technical replicates. **(H)** CD19 transfer to CAR T cells following a 30-minute coculture of FMC63 CAR or CARCSTA T cells with K562 cells expressing CD19-cLuc at a 0.5:1 effector-target ratio as determined by flow cytometry. Data is representative of at least three independent experiments using cells produced from three healthy donors. **(I)** BCMA-GFP transfer to CAR T cells following a 2-hour coculture of BCMA CAR or CARCSTA T cells and K562 cells expressing BCMA-GFP-cLuc at a 1:1 effector-target ratio as determined by flow cytometry. Data is representative of three independent experiments. **(J)** GFP transfer to CAR T cells following a 1-hour coculture with A673 cells engineered to express a CD19-GFP fusion protein at a 1:1 effector target ratio. **(K)** CD19 loss on tumor cells following a 30-minute coculture of FMC63 CAR or CARCSTA T cells and K562 cells expressing CD19-cLuc at a 0.5:1 effector-target ratio as determined by flow cytometry. Data is representative of at least three independent experiments using cells produced from three healthy donors. **(L)** BCMA loss on U266B1 tumor cells following a 2-hour coculture of BCMA CAR or CARCSTA T cells and K562 cells expressing BCMA-GFP-cLuc at a 1:1 effector-target ratio as determined by flow cytometry. Data is representative of three independent experiments. **(M)** GFP loss on A673 tumor cells following a 1-hour coculture of FMC63 CAR or CARCSTA T cells and A673 cells expressing CD19-GFP at a 1:1 effector-target ratio as determined by flow cytometry. **(J/M)** Data is representative of two independent experiments using cells produced from three healthy donors. **(N)** Total CAR T cells following a 30-minute coculture of FMC63 CAR or CARCSTA T cells and K562 cells expressing CD19-cLuc at a 0.5:1 effector-target ratio as determined by flow cytometry. Data is representative of at least three independent experiments using cells produced from three healthy donors. **(O)** Total CAR T cells following a 2-hour coculture of BCMA CAR or CARCSTA T cells and K562 cells expressing BCMA-GFP-cLuc at a 1:1 effector-target ratio as determined by flow cytometry. Data is representative of three independent experiments. **(N/O)** CAR T cell numbers are normalized to wells containing only CAR T cells using counting beads. **(P)** CD19 transfer to CAR T cells following a 1-hour coculture of FMC63 CAR or CARCSTA T cells and primary B cell acute lymphoblastic leukemia (B-ALL) cells at a 1:1 effector-target ratio as determined by flow cytometry. **(Q)** CD19 loss on primary B-ALL cells following a 1-hour coculture of FMC63 CAR or CARCSTA T cells and primary B-ALL cells at a 1:1 effector-target ratio as determined by flow cytometry. **(R)** Total CAR T cells following a 1-hour coculture of FMC63 CAR or CARCSTA T cells and primary B-ALL cells at a 1:1 effector-target ratio as determined by flow cytometry. **(Q-R)** Data is representative of three independent experiments. **(H-R)** Statistical significance was determined by one-way ANOVA.

We hypothesized that this effect on CAR T cell survival in short-term co-cultures could also lead to increased long-term CAR T cell persistence and anti-tumor activity of CAR_CSTA_ T cells. We therefore next performed a serial cytotoxicity assay to determine CAR_CSTA_ T cell anti-tumor activity over an extended period of time (Fig. 6A). We found that CAR_CSTA_ T cells significantly prolonged tumor control (Fig. 6B) and observed increased CAR T cell numbers over time (Fig. 6C). In an *in vivo* B-ALL tumor model (Fig. 6D), we observed tumor control in all groups treated with CD19-targeting CAR T cells (Fig. 6E, Suppl. Fig. S7A) and again found significantly increased CAR_CSTA_ T cell numbers (Fig. 6F). When analyzing the phenotype of these persisting CAR T cells, we found that CSTA overexpression did not significantly alter T cell subset composition (Fig. 6G). We next explored expression of exhaustion markers in CAR T cells from *in vitro* serial cytotoxicity and *in vivo* experiments. We found that, despite their substantially increased persistence in both experiments, CAR_CSTA_ cells showed significantly increased levels of exhaustion (Fig. 6H, Suppl. Fig. S7B). To explore potential causes of this increased exhaustion, we next performed bulk RNA sequencing of FMC63 CAR and CAR_CSTA_ T cells at the end of production. We found that CSTA overexpression resulted in the spontaneous upregulation of Cytokine-Inducible SH2 containing protein (CISH), which has previously been implicated as a key driver of CAR T cell exhaustion (Fig. 6I). To determine if this upregulation of CISH could have caused the increased exhaustion of CAR_CSTA_ T cells, we next established an efficient CRISPR/Cas9-mediated CISH knockout (Suppl. Fig. S7C/D). CISH^KO^ did not result in changes to T cell subset composition at the end of production (Suppl. Fig. S7E) and did not affect inhibition of CMT by CSTA overexpression (Suppl. Fig. S7F-H) indicating that the effect of CSTA on CMT is not mediated by CISH. We next performed a serial cytotoxicity assay using CAR_CSTA_ cells with and without CISH^KO^ (Fig. 6J) and found that CISH^KO^ significantly prolonged tumor control and further increased CAR T cell persistence over time (Fig. 6K/L). In addition, CISH^KO^ almost entirely prevented exhaustion in CSTA-expressing CAR T cells in vitro (Fig. 6M/N), indicating that CSTA-induced upregulation of CISH caused the increased levels of exhaustion in CAR_CSTA_ cells. We validated this finding in the same *in vivo* model described above and again observed complete tumor control and increased persistence in CAR_CSTA_ CISH^KO^ CAR T cells (Fig. 6O, Suppl. Fig. S7I). We next hypothesized that CMT could be more pronounced in the solid tumor setting due to the higher density of tumor cells leading to more extensive antigen transfer and fratricide. We therefore established a solid tumor *in vivo* model by intratibial implantation of A673 Ewing sarcoma cells engineered to express CD19 (Fig. 6P). This model was characterized by rapid localized tumor growth and efficient tumor control by systemically injected FMC63 and CAR_CSTA_ CISH^KO^ CAR T cells (Fig. 6Q, Suppl. Fig. S7J). In this model, we found that CSTA expression did not alter CAR T cell numbers in the peripheral blood (Fig. 6R) but substantially increased intratumoral CAR_CSTA_ T cell numbers indicating that CSTA overexpression does not negatively affect tumor infiltration by CAR T cells (Fig. 6S) and instead efficiently prevents intratumoral CAR T cell fratricide. Exhaustion and phenotype of CAR_CSTA_ CISH^KO^ cells was comparable to FMC63 CAR T cells indicating that CISH^KO^ robustly prevents CSTA-mediated exhaustion in solid and hematologic settings (Fig. 6T/U, Suppl. Fig. S7K).

**Figure 6:**
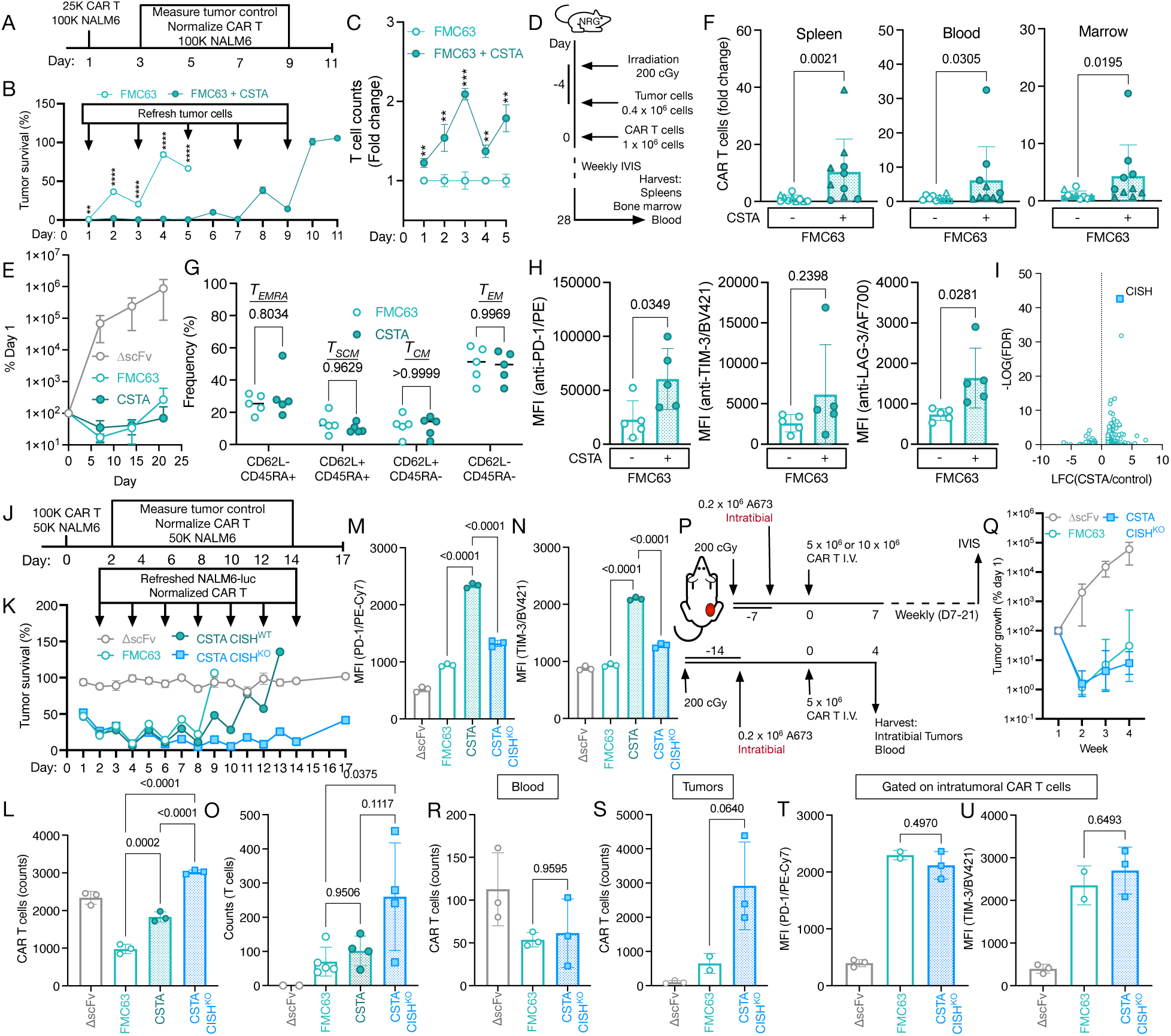
Cystatin A overexpression improves long-term CAR T cell persistence. **(A)** Schema of *in vitro* experiment to determine long-term CAR T cell persistence and antitumor activity. **(B)** Survival of CD19-GFP-expressing NALM6-Fluc cells at a 0.5:1 effector-target ratio using a luminescence-based cytotoxicity assay. Data indicates mean ± S.D. from three technical replicates. Tumor survival was normalized to untreated tumor cells. Data is representative of two independent experiments. Statistical significance comparing FMC63 CAR and CARCSTA T cells was determined by multiple two-tailed Student’s *t*-tests**. (C)** FMC63 CAR and CARCSTA T cell counts on D1-5 of the *in vitro* serial coculture described in Fig. 6A. Data represent mean ± S.D. of 3 technical replicates and is representative of two independent experiments. Statistical significance comparing FMC63 CAR and CARCSTA T cells was determined by multiple two-tailed Student’s *t*-tests**. (D)** Schema of *in vivo* experiment to measure NALM6 tumor control and CAR T cell persistence. **(E)** Tumor burden in mice bearing systemic NALM6-Fluc tumors and treated with ΔscFv or FMC63 CAR or CARCSTA T cells as determined by *in vivo imaging system (IVIS).* Data represent mean ± S.D. of 5 mice per group. Statistical significance was determined by two-way ANOVA. **(F)** Quantification of CAR T cells in murine spleen, peripheral blood, and bone marrow as determined by flow cytometry. Data represent mean ± S.D. of 10 mice pooled from two independent experiments, with experiments denoted by symbol. Data were normalized to the average of the respective experiment’s CSTA^−^ condition and are expressed as fold change. Statistical significance was determined by two-tailed Student’s *t*-test. **(G)** Phenotype of FMC63 CAR/CARCSTA T cells isolated from murine blood as determined by flow cytometry. Statistical significance was determined by two-way ANOVA. **(H)** PD-1, TIM-3, and LAG-3 expression on FMC63 CAR or CARCSTA T cells isolated from murine blood as determined by flow cytometry. Statistical significance was determined by two-tailed Student’s *t*-test. **(G/H)** Data represent mean ± S.D. of 5 mice per group. **(I)** Differential mRNA expression in sorted FMC63 CAR and CARCSTA T cells after manufacturing as determined by bulk RNA sequencing. Data is representative of cells from two independent productions. **(J)** Schema of *in vitro* experiment to determine long-term CAR T cell persistence and antitumor activity. **(K)** Survival of NALM6-Fluc cells at a 2:1 effector target ratio using a luminescence-based serial cytotoxicity assay. On Days 2, 4, 6, 8, 10, 12, and 14 CAR T cells were normalized and fresh tumor cells were added. Statistical significance was determined by two-way ANOVA. **(L)** Total CAR T cells on Day 10 of the *in vitro* serial coculture described in Fig. 6J. **(M)** PD-1 expression on CAR T cells on Day 5 of *in vitro* serial coculture described in Fig. 6J. **(N)** TIM-3 expression on CAR T cells on Day 5 of *in vitro* serial coculture described in Fig. 6J. **(K-N)** Data represent mean ± S.D. of three technical replicates and is representative of two independent experiments. **(O)** Quantification of CAR T cells in bone marrow 28 days after intravenous treatment with the indicated CAR T cell conditions as determined by flow cytometry. Data represent mean ± S.D. of 2-5 mice per group. Statistical significance was determined by one-way ANOVA. **(P)** Schema of *in vivo* experiment to measure solid tumor control and CAR T cell infiltration into solid tumors. **(Q)** Tumor burden in mice bearing intratibial CD19-expressing A673-Fluc tumors and treated intravenously with ΔscFv or FMC63 CAR CISH^WT^ or CARCSTA CISH^KO^ T cells as determined by *in vivo imaging system (IVIS).* Data represent mean ± S.D. of 3-4 mice per group. **(R)** Quantification of CAR T cells in peripheral blood four days after treatment with the indicated CAR T cell conditions as determined by flow cytometry (n = 2-3 per group). Statistical significance was determined by one-way ANOVA. **(S)** Quantification of CAR T cells in intratibial A673 tumors four days after treatment with the indicated CAR T cell conditions as determined by flow cytometry (n = 2-3 per group). Statistical significance was determined by one-way ANOVA. **(T)** PD-1 and **(U)** TIM-3 levels on intratumoral CAR T cells four days after CAR T cell injection as determined by flow cytometry. Statistical significance was determined by one-way ANOVA. **(R-U)** Data represent mean ± S.D. of 2-3 mice per group.

Taken together, these data indicate that CSTA overexpression leads to significantly reduced CMT, as well as increased long-term CAR T cell persistence and tumor control. CSTA-mediated induction of exhaustion via CISH could be overcome by simultaneous CISH^KO^ further enhancing functional CAR T cell persistence.

## Discussion

CAR T cells have revolutionized cancer immunotherapy, but most patients receiving CAR T cell therapy relapse within 5 years of treatment^12,57,58^. Several mechanisms driving relapse after CAR T cell therapy have been described, including CAR T cell-mediated trogocytosis (CMT). CMT, the extraction of tumor antigen from the tumor cell and its subsequent incorporation into the CAR T cell membrane, is modulated by CAR affinity for target antigen^23^, actin rearrangement^22,39^, cholesterol metabolism^59^, and the CAR transmembrane domain^60^. CMT has previously been shown to result in the emergence of antigen-negative tumor cells^22–24^ and had been hypothesized to cause CAR T cell fratricide as well as CAR T cell exhaustion through persistent cis signaling^61^. Previous studies have demonstrated the presence of tumor antigen on CAR T cells following co-culture with tumor cells and that the presence of trogocytosed antigen correlates with increased apoptotic markers and exhaustion^22–24^. However, it remained unknown if this T cell dysfunction was indeed caused by CMT. In this study, we sought to determine if the presence of trogocytosed antigen causes CAR T cell dysfunction and to uncover molecular modulators of CMT to improve our understanding of the trogocytic process.

In this study, we demonstrate that CAR T cell fratricide and exhaustion can be reversed through selective degradation of trogocytosed antigen on the CAR T cells, indicating that trogocytosed antigen is indeed directly causing CAR T cell dysfunction. We show that this trogocytosed antigen degrader (TAD) approach is feasible when targeting an antigen-GFP fusion protein as well as the unmodified endogenous antigen (CD19) by using endogenous protein domains binding to the target antigen’s intracellular domain. The TAD protein targeting endogenous CD19 (TAD_CD19_) relies on the established high-affinity interaction of the Lyn kinase SH2 domain with the intracellular domain of CD19^36^. However, SH2 domains can be involved in multiple signaling pathways with their activity being highly context-dependent^62–64^. It is therefore possible that TAD_CD19_ could bind to proteins other than CD19. Because of this possibility, we mainly relied on the use of TAD_GFP_, which relies on a validated protein-protein interaction and is unlikely to result in similar off-target effects^32^. If TAD-mediated targeting of an endogenous antigen is desired in the absence of target-specific endogenous binding domains, protein binders could be generated de novo, analogous to the GFP-specific VHH domain used in this study. Future work focusing on TAD-based approaches using endogenous binding domains will have to carefully consider potential off-target reactivity.

In this study, we found that the cysteine protease cathepsin B (CTSB) produced by CAR T cells is essential for CMT but dispensable for tumor cell killing. We demonstrate that CTSB produced by CAR T cells rapidly localizes to the immune synapse upon antigen stimulation and that extracellular CTSB confers antigen extraction from tumor cells. A role of extracellular CTSB in the processing of cellular material is in line with prior studies describing the involvement of tumor-derived CTSB in the degradation and remodeling of components of the extracellular matrix, enabling tumor metastasis^45,47,48^. We further show that CTSB activity in CAR T cells is tightly regulated through the abundance of cystatins, a class of endogenous protease inhibitors^49,51^. We found that overexpression of both cystatin A and cystatin B potently inhibited CTSB in CAR T cells. While the small molecule inhibitors used throughout this study have been shown to be specific for CTSB^65,66^, the more robust inhibition of CMT by cystatins, which are able to inhibit a range of cathepsins, including cathepsin L and H, could indicate a role of other cathepsins in CMT^49,51^.

In principle, constitutive overexpression of CSTA to limit CMT could cause toxicity through systemic inhibition of CTSB or other proteases^67^. However, it has been shown previously that CTSB^KO^ mice are viable and appear indistinguishable from CTSB^WT^ mice^52–55^. In addition, we found that mice treated with CAR_CSTA_ T cells exhibit no overt signs of toxicity. While constitutive overexpression of CSTA appears to be relatively safe, additional work will need to be performed in relevant model systems to confirm our initial observations. In case toxicities are observed, the safety of CSTA overexpression could be increased by using a conditional expression system that initiates CSTA expression upon CAR target recognition or recognition of a separate tissue-specific antigen^68,69^.

In this study, we specifically focused on the role of CAR T cell-derived factors on CMT. However, CTSB is expressed by both T cells and tumor cells. Using the covalent small molecule CTSB inhibitor Ca-074-Me, we show that specific inhibition on CAR T cell-derived CTSB directly causes CMT. However it is possible that tumor-derived CTSB also contributes to this process and further work is needed to comprehensively explore this possibility.

Increased levels of cystatins other than CSTA, such as cystatin F, a type II cystatin and inhibitor of cathepsins C, L, and H, have been found to decrease cytotoxicity in NK cells and T cells^70,71^, raising the question of how CSTA overexpression may affect CAR T cell function and tumor control. In this study, we observed no differences in short-term *in vitro* cytotoxicity and cytokine secretion between FMC63 CAR and CAR_CSTA_ cells. However, we observed substantially increased long-term tumor control by CAR_CSTA_ cells *in vitro*, which was also associated with significantly increased CAR T cell numbers in hematologic and solid tumor models *in vitro* and *in vivo*. Increased tumor control by CAR_CSTA_ cells is in line with previous studies which showed that inhibiting CMT using inhibitory CARs^24^, combinatorial targeting^22^, or altering cholesterol metabolism^72^ can increase anti-tumor activity. However, in long-term *in vitro* and *in vivo* experiments, we observed significant upregulation of exhaustion markers on CAR_CSTA_ T cells. Analyzing the pre-infusion product, we found that CAR_CSTA_ T cells showed spontaneous upregulation of cytokine-inducible SH2-containing protein (CISH), a known negative regulator of T cell activation^73,74^. By including a CISH knockout step during CAR_CSTA_ production, we were able to show that the CSTA-induced exhaustion was mediated by CISH, which is in line with recent studies showing that CISH can be a major driver of CAR T cell exhaustion in some contexts ^75,76^ . Our work serves as an initial proof-of-concept showing that protease inhibition is an effective approach to limit CMT but additional work will be needed to better understand how CSTA alters CISH expression and whether it is possible to further separate the beneficial effects of CSTA overexpression from those conferring increased exhaustion.

In addition to investigating the effects of CMT on CAR T cell persistence and function, we show here that antigen density is substantially lowered on tumor cells through CMT and that CMT is detectable in patients treated with CAR T cells. However, it remains unclear how much transient antigen loss observed after CMT contributes to clinical antigen escape. Monitoring CMT with high temporal resolution *in vivo* through continuous sampling could help to answer this question by elucidating the extent of CMT-mediated antigen loss. In addition, high-resolution imaging of *in vitro* cocultures with tumor cells and CAR T cells may allow visualization of tumor cells following CMT to determine if antigen loss protects these cells temporarily from CAR T cell-mediated killing.

Trogocytosis is a highly variable phenomenon and can be affected by several factors including tumor antigen density^22^, target affinity^23^, CAR structure^22,60^, and target cell type. We observed substantial variability in CMT when analyzing the peripheral blood of patients receiving CAR T cell therapy, and recent work also shows variability of CMT in mice^60^. It remains an open question why CMT varies between patients and larger clinical immunomonitoring trials will be needed to conclusively address this question.

Our study primarily focused on approaches targeting the two clinically validated CAR T cell antigens CD19 and BCMA approved for the treatment of several hematologic malignancies; however, we also show that CMT can be observed in solid malignancies when targeting different, previously validated CAR antigens. While CMT appears to occur across CAR target antigens, it remains an open question whether transfer of functionally essential and/or tumorigenic proteins, the loss of which would result in rapid target cell death, would be equally susceptible to this process. In the solid tumor setting, exhaustion-driven T cell dysfunction is generally more pronounced^77,78^ and CMT may play a key role due to a denser tumor microenvironment and limited T cell mobility, which may favor more extensive intratumoral fratricide. High-resolution intratumoral imaging may provide critical insights into CMT dynamics to inform future strategies to reduce the impact of this process.

In conclusion, we show that CMT directly causes CAR-mediated fratricide and exhaustion and that CTSB is essential for CMT. CTSB activity is controlled by the abundance of endogenous human cystatins and cystatin overexpression results in significantly reduced CMT, fratricide, exhaustion, and increased CAR T cell persistence and anti-tumor activity. CSTA overexpression is an effective approach to limit the negative consequences of CMT but requires additional engineering, such as CISH knockout, to prevent CSTA-mediated CAR T cell exhaustion. Our findings provide evidence that CMT is mechanistically distinct from cytotoxic function in CAR T cells and support the rational targeting of this resistance mechanism in CAR T cells.

## MATERIALS AND METHODS

### Study approval

All recombinant DNA and biosafety work was approved by the institutional biosafety committees at University of Maryland, Baltimore (protocol IBC-6040). Animal experiments were approved by the institutional animal care and use committee at the University of Utah (protocol 18-1104).

### Cell lines and primary human cells

Raji, NALM6, Daudi, DB, Toledo, MM.1S, RPMI8226, U266B1, K562, A673, TC-71, and Phoenix-AMPHO cells were purchased from the American Type Culture Collection (ATCC) and cultured according to ATCC instructions. Lenti-X 293T cells were purchased from Takara and cultured according to the manufacturer’s instructions. Cell lines were authenticated by their respective supplier. Healthy donor buffy coats were obtained from the New York Blood Center. PBMCs from healthy donors were isolated from buffy coats by density gradient using FillPaque (GE) as previously described^27,30^. Primary human T cells were cultured in AIM-V medium (Invitrogen 12055-083) supplemented with 5% Human serum (Sigma H4522-100ML), 1% Pen/Strep (Thermofisher 15140-122) (T cell media), and 40 IU IL-2 (R&D Systems #202-IL-10). All cells were cultured at 37°C, 5% CO2.

### Vector constructs

All vectors generated for this study were produced by Twist Biosciences. FMC63-nLuc contains the CD19-specific scFv fragment FMC63 ^79^. All CAR constructs contain the CD8α hinge/transmembrane domain, 4-1BB costimulatory domain, and CD3z domain.

For some constructs, the full-length sequence of human cystatin A (UniProt, P01040) was synthesized and cloned downstream of the respective CAR and nLuc fragment separated by a P2A sequence (Twist Bioscience). DNA was isolated using Endofree Plasmid Maxi Kit (Qiagen 12362) following the manufacturer’s protocol. Plasmid concentration was measured using a NanoDrop One instrument (Thermo). All DNA constructs were stored at -20°C.

### Clinical data

Whole blood was drawn from patients receiving CAR T cell therapy 7-28 days after CAR T cell injection. Peripheral blood mononuclear cells (PBMCs) were isolated by density gradient using Ficoll Paque as previously described^27,30^. PBMCs were stained with anti-hCD3, anti-hCD4, anti-hCD8, anti-CAR (Miltenyi #130-127-342), and anti-hCD19, anti-hCD27, anti-hCD137, and 7-AAD and analyzed by flow cytometry. Samples were collected under Institutional Review Board (IRB)-approved protocol 2043GCCC (IRB H0091736, PI D. Atanackovic).

### Gamma retrovirus Production

Gamma retrovirus was produced using Phoenix-AMPHO cells (ATCC, catalog no. CRL-3213). Phoenix-AMPHO cells were transiently transfected with 16 µg of plasmid DNA using Opti-MEM Reduced Serum Medium (Thermofisher, catalog no. 31985070) and lipofectamine 2000 (Invitrogen 11668-019) according to manufacturer’s instructions. During transfection, cells were cultured in Dulbecco’s Modified Eagle Medium (DMEM, Thermofisher 11995073) supplemented with 10% FBS. Virus-containing supernatant was filtered using Steriflip™ Sterile Disposable Vacuum Filter Units (Millipore Sigma SEIM003M00). Virus was concentrated using Retro-X Concentrator (Takara, 631456). The following day, concentrated virus was centrifuged at 1500 x g for 45 minutes at 4°C. Supernatant was removed and concentrated virus was resuspended in 1.5 mL complete T cell media.

### Transgenic T cell production and expansion

CAR T cells were generated as previously described^23,27,30^. Buffy coats from three healthy donors were obtained from the New York Blood Center and peripheral blood mononuclear cells (PBMCs) were isolated using Ficoll-Paque and cryopreserved until use. PBMCs were thawed and cultured overnight in complete T cell media. PBMCs were stimulated for 2 days with anti-CD3/anti-CD28 T cell activation beads (Thermo, catalog no. 11131D) in the presence of interleukin-2 (IL-2; 40 IU/ml; R&D Systems, catalog no. 202-IL-010) in complete T cell media and incubated at 37°C, 5% CO_2_. Bead-stimulated cells were transferred to RetroNectin-coated (Takara) virus–containing plates and incubated overnight. Transduction was repeated the next day before counting and diluting cells to 0.4 × 10^6^ cells/ml. After the second transduction, cells were grown for an additional 7 days before removing beads using a DynaMag-15 magnet (Thermo Fisher Scientific). IL-2 was replenished every 2 days to 40 IU/ml. Cells were frozen in 90% fetal bovine serum/10% dimethyl sulfoxide and stored in liquid nitrogen. CAR T cell transduction efficiency and phenotype were determined by flow cytometry. CAR T cells were washed with FACS buffer and stained with antibodies targeting hCD3, HA, hCD4, hCD8, hCD95, hCD62L, and hCD45RA. Because BCMA CAR_CSTA_ T cells showed substantially lower transduction efficiency than conventional BCMA CAR T cells, after production, we FACS sorted both products following anti-HA (CAR) staining using a FACSAria cell sorter (BD) prior to subsequent functional assays.

To generate CISH^KO^ CAR T cells, on day 5, anti-CD3/CD28 T Activator beads (Thermo Fisher) were removed from the transduced cells by magnetic separation. CRISPR ribonucleoprotein (RNP) particles containing 25 *μ* g Cas9 only (mock/WT) or Cas9 and 150 pmol synthetic TrueGuide gRNA (CISH target sequence: 5’-GACAGCGUGAACAGGUAGCU-3’) were prepared according to the manufacturer’s instructions (Thermo). 3 x 10^6^ cells were electroporated in the presence of CRISPR-RNPs using the NEON electroporation system (Thermo) using the following settings: 1,600 V, 10 ms, 3 pulses. Following the electroporation, cells were transferred to prewarmed media and incubated for 2 hours at 37°C. Following the incubation, new anti-CD3/CD28 T cell activation beads were added to the electroporated cells at a ratio of 3:1, and cells were grown for an additional 6 days before removing beads using a DynaMag-15 magnet (Thermo Fisher). IL-2 was replenished every 2 days to 40 IU/ml. Cells were frozen in 90% FCS/10% DMSO and stored in liquid nitrogen.

### Imaging Substrate Preparation

Eight-well chambers (Cellvis, catalog no. C8-1.5H-N) were used for all experiments. For CAR T cell activation on CD19-coated surfaces, 8-well chambers were coated with 0.01% poly-L-lysine (PLL) solution diluted in distilled water for 10 minutes at room temperature. PLL was aspirated from each well, and the chambers were allowed to air-dry for 1 hour at 37°C. PLL-coated dishes were then incubated overnight at 4°C with a 10 μg/mL solution of NeutrAvidin (Thermo Scientific, catalog no. 31000) in 1x Dulbecco’s phosphate-buffered saline (DPBS). After overnight incubation, coated wells were washed with 1x DPBS at room temperature and incubated at 37°C with a 10 μg/mL solution of biotinylated human CD19 protein (ACRO Biosystems catalog no. CD9-H82E9) in DPBS for 2 hours at 37°C. Prior to the experiment, coated wells were washed three times with RPMI 1640 phenol red-free imaging medium. For coculture experiments, 8-well chambers wells were incubated with 10 μg/mL fibronectin (MilliporeSigma, catalog no. 34-163-11MG) in DPBS at room temperature for 1 hour prior to seeding with 293T cells stably expressing CD19-GFP. 293T cells were seeded at a concentration of 5 x 10^5^ cells per well in complete growth media, followed by an overnight incubation at 37°C, 5% CO_2_. Prior to imaging, wells were washed three times with warm complete imaging media, consisting of a 1:1 ratio of RPMI 1640 supplemented with 5% fetal bovine serum (FBS) and DMEM supplemented with 10% FBS. The washing process was performed thrice to ensure removal of any residual media while leaving a known volume in the wells.

### Confocal Microscopy

Confocal microscopy was conducted using an inverted microscope (Nikon Ti-E PFS, Nikon Inc.) equipped with a 100× Silicone objective lens. Imaging was performed with a Prime BSI camera (Photometrics). Image acquisition protocols were managed using Nikon Elements software, and images were cropped in FIJI for further analysis. All live cell imaging was done with imaging chambers placed in a stage-top Okolab Incubator (Okolab S. R. L.) pre-equilibrated to 37°C with 5% CO_2_. For live cell imaging on glass, activated CAR T cells suspended in RPMI 1640 medium supplemented with 5% FBS were deposited onto CD19-coated surfaces. Imaging was started between 3 to 6 minutes after CAR T cells expressing CTSB-mCherry were added to a biotin-CD19-coated coverslip. For each well, timelapse images of a 3D volume (planes with z spacing of 0.3 μm to span the cell from the top to the bottom) were acquired every 3 minutes for 60 minutes. For coculture experiments, imaging was started between 3 to 6 minutes after CAR T cells expressing CTSB-mCherry were dropped onto a layer of 293T cells expressing CD19-GFP seeded on a coverslip, at a concentration of 7 x 10^4^ cells per drop. Brightfield imaging was used to identify cells attached to the apical surface of HEK293 cells. The synaptic plane between a CAR-T cell and a HEK293 cell was identified and designated as the home plane for acquisition of Z-stack time-lapse movies. Two-channel images using 488 nm and 561 nm lasers (for GFP and mCherry imaging respectively) were acquired every 3 minutes for 1 hour and 15 minutes utilizing a Z-spacing of 0.3 or 0.6 μm, with brightfield images taken at the home plane.

Image analysis was carried out in MATLAB (Mathworks, Inc.) using custom scripts. The plane of the synapse was determined using the actin channel. After background subtraction, axial intensity gradients are estimated for every voxel of sufficient intensity within the ROI. Below the cell, these gradients are typically positive due to the Airy pattern of the PSF. The synapse is taken as the first plane for which the gradients are no longer consistently positive.

### Estimating the average distance of CTSB to the synapse

The center of fluorescence (COF) of CTSB is defined similarly to the center of mass. The voxel positions are weighted by CTSB intensity after background subtraction and thresholding to obtain the COF. The average CTSB distance is then calculated from the axial (z) distance of the Cathepsin B COF to the plane of the synapse.

### Characterizing CTSB axial dispersion

The axial dispersion is an estimate of the average distance of CTSB molecules to the COF. The axial (z) distance of each voxel to the COF is determined, and the axial dispersion is defined as the average of these distances weighted by CTSB intensity after background subtraction and thresholding. Voxels with sufficiently low signal do not contribute to the calculation due to the thresholding procedure.

### CTSB clustering at the synapse

The pair auto-correlation function g(r) of CTSB is computed at the synapse using the actin channel as a mask^80^. The actin channel is segmented by smoothing with a Gaussian filter, generating an initial mask by applying k-means clustering (k = 2) after a log transformation, and then applying morphological operations to connect and smooth the initial mask. The clustering coefficient, g_ave_, is computed by averaging over all radial bins with 0.25 m or 0.5 m.

### CompLuc-based trogocytosis assay

Transduced, cryopreserved nLuc+ CAR T cells were thawed and cultured in complete T cell media supplemented with 40 IU IL-2 for 48 hours prior to use. nLuc-expressing CAR T cells were cocultured with CD19-cLuc-expressing K562 tumor cells at the specified effector-target ratios. CAR T cells and tumor cells were resuspended in Opti-MEM reduced serum media. Live Cell Substrate (Promega, catalog no. N2011) was prepared according to manufacturer’s instructions. K562 tumor cells and prepared Live Cell Substrate were added to wells of a black 96-well plate and luminescence was measured to assess baseline luminescence. CAR T cells were added to appropriate wells and luminescence was measured every minute for three hours at 37°C. Luminescence was measured using a Spark multi-mode plate reader (Tecan).

### Flow cytometry-based trogocytosis assay

A flow-cytometry based trogocytosis assay was used to confirm results observed in CompLuc, to assess CD19 or BCMA levels on CAR T cells and tumor cells, and to quantify CAR T cells. 5 x 10^4^ target cells were seeded in wells of a 96-well round bottom plate. Various ratios of CAR T cells produced from one of three healthy donors were cocultured with target cells for 1 hour at 37°C, 5% CO_2_. Following coculture, cells were resuspended by gentle pipetting and transferred to wells of a 96-well V bottom plate for washing and staining. Cells were stained with Zombie violet or Zombie NIR fixable viability dye, and antibodies for hCD3, HA, hCD19 or BCMA. When assessing exhaustion, cells were stained with antibodies targeting PD-1, LAG-3, and TIM-3. When assessing phenotype, cells were stained with antibodies targeting CD45RA and CD62L. Accucheck counting beads (Life Technologies) were added to the cells for normalization. Samples were acquired on an LSR II flow cytometer (BD) or an Aurora full-spectrum flow cytometer (Cytek).

### Luciferase-based cytotoxicity assay

To determine *in vitro* CAR T cell cytotoxicity, cell lines (Raji, NALM6, Daudi, Toledo MM.1S, RPMI8226, U266B1) were transduced with pHIV-Luc-ZsGreen lentivirus and sorted on a FACS Aria flow cytometer (BD) for GFP expression. 3 x 10^4^ target cells were seeded in wells of a 96-well round bottom plate. CAR T cells from one of three healthy donors were cocultured with target cells at the indicated effector-target ratios and incubated for 16 hours at 37°C, 5% CO_2_. Following incubation, 80 μL of supernatant was harvested from each well. Cells were suspended by gentle pipetting, and 100 μL was transferred to a 96-well black flat bottom plate. D-Luciferin (Gold Biotechnology, catalog no. LUCNA-2G) at 150 μg/mL was added to the cells and incubated for 5 minutes at 37°C. Luminescence was determined on a Spark multimode plate reader (Tecan).

### Serial coculture repeat stimulation assay

To determine the long-term *in vitro* control and exhaustion of CAR T cells, luciferase-expressing tumor cells were plated at 5 x 10^4^ cells/well. CAR T cells were cocultured at a defined effector-target ratio and incubated for 24-48 hours at 37°C, 5% CO_2_. Following incubation, cells were transferred to a 96-well black flat bottom plate. D-luciferin was added to cells and luminescence was determined on a Spark multimode plate reader (Tecan). After measuring luminescence, cells were washed with FACS buffer and stained with Zombie NIR fixable viability dye (Biolegend) and antibodies targeting CD19, HA, PD-1, TIM-3, CD45RA, and CD62L. Accucheck counting beads (Life Technologies) were added to the cells for normalization. Next, remaining CAR cells were pooled together and normalized based on expansion. CAR T cells were redistributed to wells and fresh tumor cells were added to each well. Plates were incubated for 48 hours at 37°C, 5% CO_2_. Luminescence measurements and normalization were repeated until a loss of cytotoxicity was observed.

### Cystatin A enzyme-linked immunosorbent assay

Cystatin A concentration was assessed using a Human Cystatin A ELISA kit (Invitrogen, catalog no. EH140RB). Total cell lysates were extracted from CAR T cells using radioimmunoprecipitation assay (RIPA) buffer (Thermo) containing protease inhibitor cocktail (Roche). Total protein concentration was determined using Pierce BCA assay (Thermo). Cystatin A levels were determined by enzyme-linked immunosorbent assay (ELISA) according to manufacturer’s instructions (Invitrogen) and calculated using standard curve interpolation. Absorbance was measured at the recommended wavelength on a Spark multimode plate reader (Tecan).

### Cathepsin B activity assay

Cathepsin B activity was assessed using the InnoZyme Cathepsin B Activity Assay Kit (MilliPore Sigma, catalog no. CBA001). Total cell lysates were extracted from CAR T cells using the provided cell lysis buffer according to manufacturer’s instructions. Total protein concentration was determined using Pierce BCA assay (Thermo Fisher Scientific). Cathepsin B activity was determined fluorometrically according to manufacturer’s instructions (Calbiochem). Fluorescence was measured on a Spark multimode plate reader (Tecan).

### Treatment of CAR T cells with inhibitors

FMC63 CAR T cells were treated with small-molecule inhibitors targeting Actin (inhibitor: Cytochalasin D, Sigma Aldrich catalog no. C8273), Dynamin (inhibitor: DynaSore, Sigma Aldrich catalog no. D7693), Cathepsin B (inhibitor: Ca-074-Me, SelleckChem catalog no. S7420), Clathrin (inhibitor: PitStop, Abcam catalog no. ab120687), or LFA-1 (inhibitor: BI-1950, Boehringer Ingelheim) at the indicated concentrations for one hour. Following treatment, cells were washed with T cell media and centrifuged at 400 x g for 5 minutes. To determine the effect of the non-membrane permeable CTSB inhibitor CA-074 (Millipore Sigma, catalog no. 205530), the inhibitor was added directly to the co-culture at the indicated concentrations. As a control, CAR T cells treated with DMSO. CAR T cells were cocultured with CD19-expressing tumor cells for the indicated intervals.

### Western blot

Total cell lysates were extracted from CAR T cells using radioimmunoprecipitation assay (RIPA) buffer (Thermo Fisher Scientific) containing protease inhibitor (Roche). Total protein concentration was determined using Pierce BCA assay (Thermo Fisher Scientific). Samples were separated by SDS–polyacrylamide gel electrophoresis, and separated proteins were transferred to nitrocellulose membranes using an iBlot2 transfer system (Thermo Fisher Scientific). Membranes were blocked with 5% nonfat milk–tris-buffered saline and incubated with primary antibodies against CD19, mCSTA, CISH, or β-actin. Membranes were washed and developed using species-specific secondary anti-IgG/horseradish peroxidase antibodies (R&D Systems) and Western Lightning Plus-ECL solution (PerkinElmer). Bands were visualized and quantified on an iBright 1500 imaging system (Thermo Fisher Scientific).

### *In vivo* cystatin A toxicity

6-8 week-old NOD.*Cg-Rag1^tm1Mom^Il2rg^tm1Wjl^*/SzJ (NRG) mice (Jackson Laboratory) were irradiated with a sublethal dose of 200 cGy (Rad-Source 2000) and that same day injected with the indicated number of CAR T cells into the lateral tail vein. To assess toxicity, animals were weighed daily and monitored for signs of distress in accordance with institutional regulations. After 14 days, mice were euthanized and lungs, spleens, liver, and gastrointestinal organs were collected, sectioned, and underwent H&E staining to assess organ structure. Slides were imaged using a BZ-X810 fluorescence microscope (Keyence) and BZ-X800 analyzer.

### Secretome analysis

CAR and CAR_CSTA_ secretomes were analyzed by Codeplex assay (IsoPlexis). Following production and cryopreservation, CAR and CAR_CSTA_ T cells produced from three healthy donors were thawed and cultured in T cell media + 40IU IL-2 for 48 hours. Cells were activated using anti-CD3/CD28 activation beads (Thermo Fisher) and incubated for 24 hours. Supernatants were collected after 24 hours, and cytokine profiling was measured using IsoPlexis Codeplex (Kcas Bio).

### Bulk RNA sequencing

Following production and cryopreservation, CAR and CAR_CSTA_ T cells from two independent productions were thawed and cultured in T cell media + 40IU IL-2 for 48 hours. After 48 hours, CAR T cells were washed with FACS buffer, stained with anti-CD3, anti-HA, and DAPI, and sorted for CAR expression. Sorted CAR T cells were then pelleted and flash frozen. RNA was extracted, polyA-enriched, and sequenced using Illumina NovaSeq S4 PE100 sequencing targeting 10 million read pairs per sample.

### *In vivo* cystatin A model of CAR T cell tumor control and persistence

6–8-week-old male NOD.*Cg-Rag1^tm1Mom^Il2rg^tm1Wjl^*/SzJ (NRG) mice (Jackson Laboratory) were irradiated with a sublethal dose of 200 cGy (Day 0). That same day, mice were injected with 4 x 10^5^ NALM6 tumor cells via tail vein injection. On Day 4, mice were injected with 1 x 10^6^ FMC63 CAR or CAR_CSTA_ T cells or CAR T cells lacking a binding domain (ΔscFv) via tail vein injection. Animals were weighed twice weekly and monitored for signs of distress in accordance with institutional regulations. Tumor burden was assessed weekly in prone and supine position by an *in vivo* imaging system (IVIS). For *in vivo* imaging, mice received an intraperitoneal injection of 3.3 mg D-luciferin (GOLDBIO # LUCK-10G). On day 28, animals were euthanized and tissues collected for analysis by flow cytometry. Average radiance values (p/s/cm²/sr) were determined using Living Image 4.8 software (PerkinElmer).

### *In vivo* model of CAR T cell solid tumor infiltration

6-8 week-old NOD.*Cg-Rag1^tm1Mom^Il2rg^tm1Wjl^*/SzJ (NRG) mice (Jackson Laboratory) were irradiated with a sublethal dose of 200 cGy (Rad-Source 2000) and that same day injected intratibially with the indicated number of luciferase-expressing A673 cells (A673-Fluc) expressing CD19-GFP. On Day 14 after tumor cell injection, the indicated number of ΔscFv, FMC63 CISH^WT^ CAR, or CISH^KO^ CAR_CSTA_ T cells were injected into the lateral tail vein. 4 days after CAR T cell injection, mice were euthanized and tumors were collected and dissociated using the Human Tumor Dissociation Kit (Miltenyi Biotec, cat # 130-095-929) and the OctoMACS tissue dissociator (Miltenyi Biotec). CAR T cells were quantified and phenotyped by flow cytometry.

### *In vivo* model of CAR T cell solid tumor control

6-8 week-old NOD.*Cg-Rag1^tm1Mom^Il2rg^tm1Wjl^*/SzJ (NRG) mice (Jackson Laboratory) were irradiated with a sublethal dose of 200 cGy (Rad-Source 2000) and that same day injected intratibially with the indicated number of luciferase-expressing A673 cells (A673-Fluc) expressing CD19-GFP. On Day 7 after tumor cell injection, the indicated number of ΔscFv, FMC63 CISH^WT^ CAR, or CISH^KO^ CAR_CSTA_ T cells were injected into the lateral tail vein. Tumor burden was assessed weekly by *in vivo* imaging system (IVIS). For *in vivo* imaging, mice received an intraperitoneal injection of 3.3 mg D-luciferin (GOLDBIO # LUCK-10G). Peripheral blood was collected on Day 19 after CAR T cell injection for CAR T cell quantification and phenotyping by flow cytometry.

### Statistical Analysis

The respective statistical tests are stated in the figure legends. Generally, statistical significance between two groups was determined by two-sided Student’s *t*-test or Mann-Whitney U test. Statistical significance between groups of three or more was determined by one-or two-way analysis of variance (ANOVA). All statistical tests were performed using Prism 10 (GraphPad). Results were considered significant when *p* < 0.05. * = p < 0.05; ** = p < 0.01; *** = p < 0.001; **** = p < 0.0001.

## Acknowledgements

The authors thank the Flow Cytometry Shared Service of the University of Maryland Marlene and Stewart Greenebaum Comprehensive Cancer Center for help with sorting cells and panel design. We thank Dr. Nevil Singh at the University of Maryland, Baltimore for use of the Keyence BZ-X810 fluorescence microscope. RNA sequencing and the associated analysis was performed by Maryland Genomics at the Institute for Genome Sciences, University of Maryland School of Medicine. Research reported in this publication utilized the Preclinical Research Shared Resource at Huntsman Cancer Institute at the University of Utah and was supported by the National Cancer Institute of the National Institutes of Health under Award Number P30CA042014. This work was supported by funds through the Maryland Department of Health’s Cigarette Restitution Fund Program (CH-649-CRF), the National Cancer Institute Cancer Center Support Grant (P30CA134274), the National Institutes of Health to AU (R35 GM145313), and by an NIAID-funded predoctoral fellowship to JMB (T32 AI095190). The plasmid pHIV-Luc-ZsGreen was a gift from Dr. B. Welm (Addgene #39196). The plasmid SFG.CNb30_opt.IRES.eGFP was a gift from Dr. M. Pule (Addgene #22493). The content is solely the responsibility of the authors and does not necessarily represent the official views of the NIH.

## Author contributions

KAD and TL conceived the project, planned and performed experiments, analyzed data, and wrote the manuscript. KAD performed the majority of the described *in vitro* experiments, generated the majority of transgenic T cells, developed the CompLuc assay, identified and validated the role of CTSB in CMT, and developed the CSTA-based approach of CMT inhibition in CAR T cells. KN and AP performed confocal microscopy and fixed coculture staining and imaging. FF wrote analysis scripts for CTSB dispersion. MGF and SVR generated FOLR⍺ CAR T cells. JMB planned experiments, analyzed data, and edited the manuscript. EG performed CAR T cell phenotyping and analyzed data. WS, ER, and DL planned and performed *in vivo* models and analyzed data. DY, RM, and AW planned experiments and analyzed data. XF developed full-spectrum flow cytometry panel and performed flow cytometry analysis. DA supervised CAR T cell phenotyping. AU planned and supervised microscopy experiments and analyzed data. TL supervised all work related to this project. All authors reviewed and approved the manuscript.

## REFERENCES

1. Sadelain, M. CD19 CAR T Cells. Cell 171, 1471 (2017).

2. Maude, S. L. et al. Tisagenlecleucel in Children and Young Adults with B-Cell Lymphoblastic Leukemia. N. Engl. J. Med. 378, 439–448 (2018).

3. Sadelain, M., Rivière, I. & Riddell, S. Therapeutic T cell engineering. Nature 545, 423–431 (2017).

4. Raje, N. et al. Anti-BCMA CAR T-Cell Therapy bb2121 in Relapsed or Refractory Multiple Myeloma. N. Engl. J. Med. 380, 1726–1737 (2019).

5. D’Agostino, M. & Raje, N. Anti-BCMA CAR T-cell therapy in multiple myeloma: can we do better? Leukemia 34, 21–34 (2020).

6. Romanski, A. et al. CD19-CAR engineered NK-92 cells are sufficient to overcome NK cell resistance in B-cell malignancies. J. Cell. Mol. Med. 20, 1287–1294 (2016).

7. Boissel, L. et al. Retargeting NK-92 cells by means of CD19-and CD20-specific chimeric antigen receptors compares favorably with antibody-dependent cellular cytotoxicity. OncoImmunology 2, e26527 (2013).

8. Cappell, K. M. & Kochenderfer, J. N. Long-term outcomes following CAR T cell therapy: what we know so far. Nat. Rev. Clin. Oncol. 20, 359–371 (2023).

9. Lee, A. Obecabtagene Autoleucel: First Approval. Mol. Diagn. Ther. 1–5 (2025) doi:10.1007/s40291-025-00771-z.

10. Purdum, A., Tieu, R., Reddy, S. R. & Broder, M. S. Direct Costs Associated with Relapsed Diffuse Large B-Cell Lymphoma Therapies. The Oncologist 24, 1229–1236 (2019).

11. Kanas, G. et al. Epidemiology of diffuse large B-cell lymphoma (DLBCL) and follicular lymphoma (FL) in the United States and Western Europe: population-level projections for 2020–2025. Leukemia & Lymphoma 63, 54–63 (2022).

12. Grover, P. et al. Chimeric antigen receptor T-cell therapy in adults with B-cell acute lymphoblastic leukemia. Blood Adv 6, 1608–1618 (2022).

13. Bethge, W. A. et al. GLA/DRST real-world outcome analysis of CAR T-cell therapies for large B-cell lymphoma in Germany. Blood 140, 349–358 (2022).

14. Pan, J. et al. Frequent occurrence of CD19-negative relapse after CD19 CAR T and consolidation therapy in 14 TP53-mutated r/r B-ALL children. Leukemia 34, 3382–3387 (2020).

15. Wudhikarn, K. et al. Interventions and outcomes of adult patients with B-ALL progressing after CD19 chimeric antigen receptor T-cell therapy. Blood 138, 531–543 (2021).

16. Finney, O. C. et al. CD19 CAR T cell product and disease attributes predict leukemia remission durability. J. Clin. Investig. 129, 2123–2132 (2019).

17. Feucht, J. et al. Calibration of CAR activation potential directs alternative T cell fates and therapeutic potency. Nat. Med. 25, 82–88 (2019).

18. L., M. S., et al. Chimeric Antigen Receptor T Cells for Sustained Remissions in Leukemia. N. Engl. J. Med. 371, 1507–1517 (2014).

19. Zhao, Z. et al. Structural Design of Engineered Costimulation Determines Tumor Rejection Kinetics and Persistence of CAR T Cells. Cancer Cell 28, 415–428 (2015).

20. Shah, N. N. & Fry, T. J. Mechanisms of resistance to CAR T cell therapy. Nat Rev Clin Oncol 16, 372–385 (2019).

21. Gardner, R. A. et al. Intent-to-treat leukemia remission by CD19 CAR T cells of defined formulation and dose in children and young adults. Blood 129, 3322–3331 (2017).

22. Hamieh, M. et al. CAR T cell trogocytosis and cooperative killing regulate tumour antigen escape. Nature 568, 112–116 (2019).

23. Olson, M. L. et al. Low-affinity CAR T cells exhibit reduced trogocytosis, preventing rapid antigen loss, and increasing CAR T cell expansion. Leukemia 36, 1943–1946 (2022).

24. Li, Y. et al. KIR-based inhibitory CARs overcome CAR-NK cell trogocytosis-mediated fratricide and tumor escape. Nat Med 28, 2133–2144 (2022).

25. Ahmed, K. A., Munegowda, M. A., Xie, Y. & Xiang, J. Intercellular Trogocytosis Plays an Important Role in Modulation of Immune Responses. Cell. Mol. Immunol. 5, 261–269 (2008).

26. Huang, J.-F. et al. TCR-Mediated Internalization of Peptide-MHC Complexes Acquired by T Cells. Science 286, 952–954 (1999).

27. Mause, E. R. V. et al. Systematic single amino acid affinity tuning of CD229 CAR T cells retains efficacy against multiple myeloma and eliminates on-target off-tumor toxicity. Sci. Transl. Med. 15, eadd7900 (2023).

28. Camviel, N. et al. Both APRIL and antibody-fragment-based CAR T cells for myeloma induce BCMA downmodulation by trogocytosis and internalization. J. Immunother. Cancer 10, e005091 (2022).

29. Cohen, A. D. et al. B cell maturation antigen-specific CAR T cells are clinically active in multiple myeloma. J. Clin. Investig. 129, 2210–2221 (2019).

30. Radhakrishnan, S. V. et al. CD229 CAR T cells eliminate multiple myeloma and tumor propagating cells without fratricide. Nat. Commun. 11, 798 (2020).

31. Bu, D.-X. et al. Pre-clinical validation of B cell maturation antigen (BCMA) as a target for T cell immunotherapy of multiple myeloma. Oncotarget 9, 25764–25780 (2018).

32. Marei, H. et al. Antibody targeting of E3 ubiquitin ligases for receptor degradation. Nature 610, 182–189 (2022).

33. Békés, M., Langley, D. R. & Crews, C. M. PROTAC targeted protein degraders: the past is prologue. Nat. Rev. Drug Discov. 21, 181–200 (2022).

34. Billerhart, M. et al. Recombinant Human CD19 in CHO-K1 Cells: Glycosylation Patterns as a Quality Attribute of High Yield Processes. Int. J. Mol. Sci. 24, 10891 (2023).

35. Heard, A. et al. Antigen glycosylation regulates efficacy of CAR T cells targeting CD19. Nat. Commun. 13, 3367 (2022).

36. Brooks, S. R., Kirkham, P. M., Freeberg, L. & Carter, R. H. Binding of Cytoplasmic Proteins to the CD19 Intracellular Domain Is High Affinity, Competitive, and Multimeric. J. Immunol. 172, 7556–7564 (2004).

37. Ruella, M. et al. Induction of resistance to chimeric antigen receptor T cell therapy by transduction of a single leukemic B cell. Nat. Med. 24, 1499–1503 (2018).

38. Dixon, A. S. et al. NanoLuc Complementation Reporter Optimized for Accurate Measurement of Protein Interactions in Cells. ACS Chem. Biol. 11, 400–408 (2016).

39. Miyake, K. et al. Trogocytosis of peptide-MHC class II complexes from dendritic cells confers antigen-presenting ability on basophils. Proc Natl Acad Sci U S A 114, 1111–1116 (2017).

40. Gilmartin, A. A., Ralston, K. S. & Petri, W. A. Inhibition of Amebic Cysteine Proteases Blocks Amebic Trogocytosis but Not Phagocytosis. J. Infect. Dis. 221, 1734–1739 (2020).

41. Reed, J., Reichelt, M. & Wetzel, S. A. Lymphocytes and Trogocytosis-Mediated Signaling. Cells 10, 1478 (2021).

42. Hudrisier, D., Aucher, A., Puaux, A.-L., Bordier, C. & Joly, E. Capture of Target Cell Membrane Components via Trogocytosis Is Triggered by a Selected Set of Surface Molecules on T or B Cells. J. Immunol. 178, 3637–3647 (2007).

43. Cavallo-Medved, D., Moin, K. & Sloane, B. Cathepsin B. AFCS Nat Mol Pages 2011, A000508 (2011).

44. Mort, J. S. & Buttle, D. J. Cathepsin B. Int. J. Biochem. Cell Biol. 29, 715–720 (1997).

45. Vidak, E., Javoršek, U., Vizovišek, M. & Turk, B. Cysteine Cathepsins and Their Extracellular Roles: Shaping the Microenvironment. Cells 8, 264 (2019).

46. Balaji, K. N., Schaschke, N., Machleidt, W., Catalfamo, M. & Henkart, P. A. Surface Cathepsin B Protects Cytotoxic Lymphocytes from Self-destruction after Degranulation. Journal of Experimental Medicine 196, 493–503 (2002).

47. Olson, O. C. & Joyce, J. A. Cysteine cathepsin proteases: regulators of cancer progression and therapeutic response. Nat Rev Cancer 15, 712–729 (2015).

48. Gabrijelcic, D. et al. Cathepsins B, H and L in human breast carcinoma. Eur J Clin Chem Clin Biochem 30, 69–74 (1992).

49. Estrada, S. et al. The Role of Gly-4 of Human Cystatin A (Stefin A) in the Binding of Target Proteinases. Characterization by Kinetic and Equilibrium Methods of the Interactions of Cystatin A Gly-4 Mutants with Papain, Cathepsin B, and Cathepsin L. Biochemistry 37, 7551–7560 (1998).

50. Pavlova, A., Krupa, J. C., Mort, J. S., Abrahamson, M. & Björk, I. Cystatin inhibition of cathepsin B requires dislocation of the proteinase occluding loop. Demonstration by release of loop anchoring through mutation of His110. FEBS Lett. 487, 156–160 (2000).

51. Pavlova, A. & Björk, I. Grafting of Features of Cystatins C or B into the N-Terminal Region or Second Binding Loop of Cystatin A (Stefin A) Substantially Enhances Inhibition of Cysteine Proteinases. Biochemistry 42, 11326–11333 (2003).

52. Guicciardi, M. E., Miyoshi, H., Bronk, S. F. & Gores, G. J. Cathepsin B Knockout Mice Are Resistant to Tumor Necrosis Factor-α-Mediated Hepatocyte Apoptosis and Liver Injury Implications for Therapeutic Applications. Am. J. Pathol. 159, 2045–2054 (2001).

53. Vasiljeva, O. et al. Reduced tumour cell proliferation and delayed development of high-grade mammary carcinomas in cathepsin B-deficient mice. Oncogene 27, 4191–4199 (2008).

54. Hook, G. et al. Cathepsin B Gene Knockout Improves Behavioral Deficits and Reduces Pathology in Models of Neurologic Disorders. Pharmacol. Rev. 74, 600–629 (2022).

55. Deussing, J. et al. Cathepsins B and D are dispensable for major histocompatibility complex class II-mediated antigen presentation. Proc. Natl. Acad. Sci. 95, 4516–4521 (1998).

56. Jenko, S. et al. Different propensity to form amyloid fibrils by two homologous proteins— Human stefins A and B: Searching for an explanation. *Proteins: Struct., Funct.*, Bioinform. 55, 417–425 (2004).

57. Xu, X. et al. Mechanisms of Relapse After CD19 CAR T-Cell Therapy for Acute Lymphoblastic Leukemia and Its Prevention and Treatment Strategies. Front. Immunol. 10, 2664 (2019).

58. Gu, T., Zhu, M., Huang, H. & Hu, Y. Relapse after CAR-T cell therapy in B-cell malignancies: challenges and future approaches. J. Zhejiang Univ.-Sci. B 23, 793–811 (2022).

59. Lu, Z. et al. ATF3 and CH25H regulate effector trogocytosis and anti-tumor activities of endogenous and immunotherapeutic cytotoxic T lymphocytes. Cell Metab. 34, 1342–1358.e7 (2022).

60. Barbera, S., et al. Trogocytosis of chimeric antigen receptors between T cells is regulated by their transmembrane domains. Sci. Immunol. 10, eado2054 (2025).

61. Ruella, M. et al. Induction of resistance to chimeric antigen receptor T cell therapy by transduction of a single leukemic B cell. Nat. Med. 24, 1499–1503 (2018).

62. Jin, L. L. et al. Tyrosine Phosphorylation of the Lyn Src Homology 2 (SH2) Domain Modulates Its Binding Affinity and Specificity*[S]. Mol. Cell. Proteom. 14, 695–706 (2015).

63. Stover, D. R., Furet, P. & Lydon, N. B. Modulation of the SH2 Binding Specificity and Kinase Activity of Src by Tyrosine Phosphorylation within Its SH2 Domain (∗). J. Biol. Chem. 271, 12481–12487 (1996).

64. Xu, Y., Harder, K. W., Huntington, N. D., Hibbs, M. L. & Tarlinton, D. M. Lyn Tyrosine Kinase Accentuating the Positive and the Negative. Immunity 22, 9–18 (2005).

65. Towatari, T. et al. Novel epoxysuccinyl peptides A selective inhibitor of cathepsin B, in vivo. FEBS Lett. 280, 311–315 (1991).

66. Murata, M. et al. Novel epoxysuccinyl peptides Selective inhibitors of cathepsin B, in vitro. FEBS Lett. 280, 307–310 (1991).

67. Turk, V. & Bode, W. The cystatins: Protein inhibitors of cysteine proteinases. FEBS Lett. 285, 213–219 (1991).

68. Smole, A. et al. Expression of inducible factors reprograms CAR-T cells for enhanced function and safety. Cancer Cell 40, 1470–1487.e7 (2022).

69. Allen, G. M. et al. Synthetic cytokine circuits that drive T cells into immune-excluded tumors. Science 378, eaba1624–eaba1624 (2022).

70. Prunk, M. et al. Extracellular Cystatin F Is Internalised by Cytotoxic T Lymphocytes and Decreases Their Cytotoxicity. Cancers 12, 3660 (2020).

71. Magister, Š., Tseng, H.-C., Bui, V. T., Kos, J. & Jewett, A. Regulation of split anergy in natural killer cells by inhibition of cathepsins C and H and cystatin F. Oncotarget 6, 22310– 22327 (2015).

72. Lu, Z. et al. ATF3 and CH25H regulate effector trogocytosis and anti-tumor activities of endogenous and immunotherapeutic cytotoxic T lymphocytes. Cell Metab. 34, 1342–1358.e7 (2022).

73. Palmer, D. C. et al. Cish actively silences TCR signaling in CD8+ T cells to maintain tumor tolerance. J. Exp. Med. 212, 2095–2113 (2015).

74. Lv, J. et al. Disruption of CISH promotes the antitumor activity of human T cells and decreases PD-1 expression levels. Mol. Ther. - Oncolytics 28, 46–58 (2023).

75. Lv, J. et al. Disruption of CISH promotes the antitumor activity of human T cells and decreases PD-1 expression levels. Mol. Ther. - Oncolytics 28, 46–58 (2023).

76. Palmer, D. C. et al. Internal checkpoint regulates T cell neoantigen reactivity and susceptibility to PD1 blockade. Med 3, 682–704.e8 (2022).

77. Newick, K., O’Brien, S., Moon, E. & Albelda, S. M. CAR T Cell Therapy for Solid Tumors. Annu. Rev. Med. 68, 139–152 (2016).

78. Sterner, R. C. & Sterner, R. M. CAR-T cell therapy: current limitations and potential strategies. Blood Cancer J. 11, 1–11 (2021).

79. Lee, D. W. et al. T cells expressing CD19 chimeric antigen receptors for acute lymphoblastic leukaemia in children and young adults: a phase 1 dose-escalation trial. Lancet 385, 517–528 (2015).

80. Veatch, S. L. et al. Correlation Functions Quantify Super-Resolution Images and Estimate Apparent Clustering Due to Over-Counting. PLoS ONE 7, e31457 (2012).

