## Supplementary Material for "Preventing trogocytosis by cathepsin B inhibition augments CAR T cell function"

*Dietze et. al.*

### SUPPLEMENTARY FIGURES

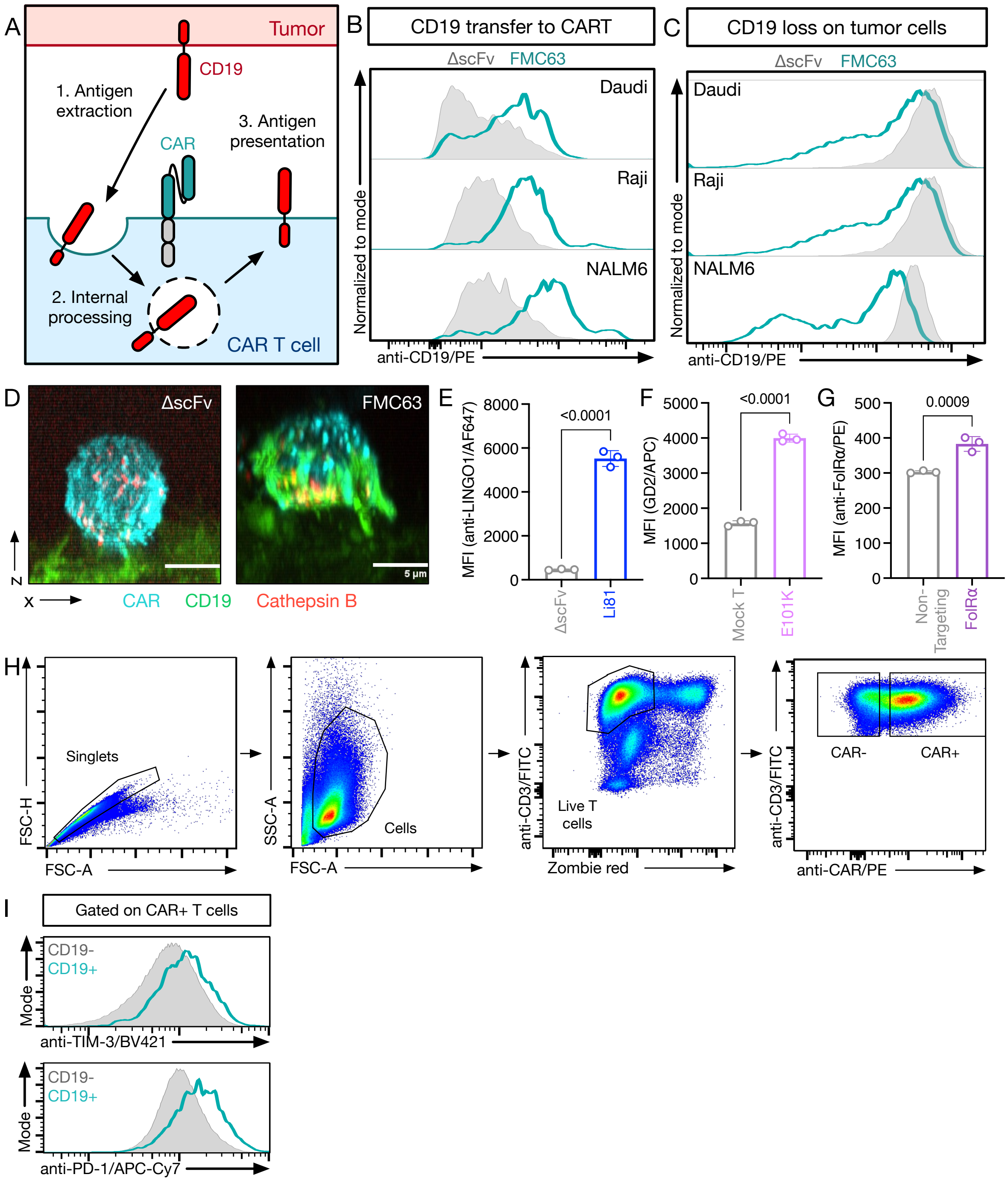


**Supplementary Figure S1: CAR T cells acquire target antigen via trogocytosis. (A)** Schema of CAR-mediated trogocytosis (CMT). **(B)** CD19 transfer to CAR T cells lacking an antigen binding domain (ΔscFv) or CD19 CAR T cells (FMC63) after a 1-hour coculture with CD19-positive cell lines at a 0.5:1 effector-target ratio using flow cytometry. **(C)** CD19 loss on tumor cells after a 1-hour coculture with ΔscFv or FMC63 CAR T cells at a 0.5:1 effector-target ratio using flow cytometry. **(B/C)** Data is representative of at least three independent experiments. **(D)** Confocal imaging z-stack showing antigen transfer from CD19-GFP-expressing 293T cells to FMC63 CAR T cells after 10 minutes. CAR is shown in cyan, CD19 is shown in green, and cathepsin B is shown in red. Scale bar represents 5$\mu$m. **(E)** LINGO1 transfer (top) to ΔscFv CAR T cells or CAR T cells targeting LINGO1 (Li81) after a 1-hour coculture with the Ewing sarcoma cell line A673 at a 1:1 effector-target ratio as determined by flow cytometry. **(F)** GD2 transfer to ΔscFv CAR T cells or CAR T cells targeting GD2 (E101K) after a 1-hour coculture with the GD2+ Ewing sarcoma cell line TC-71 at a 1:1 effector-target ratio using flow cytometry. **(E/F)** Data is representative of two independent experiments. **(G)** Folate receptor alpha (FolRα) transfer to CAR T cells targeting FolRα or an irrelevant CAR after a 1-hour coculture with the FolRα+ ovarian cancer cell line SKOV3 at a 1:1 effector-target ratio using flow cytometry. **(H)** Gating strategy for assessment of trogocytosis in clinical samples. **(I)** TIM-3 and PD-1 levels on $CD19^-$ and $CD19^+$ CAR T cells from the peripheral blood of a patient receiving CD19 CAR T cell therapy, as determined by flow cytometry.

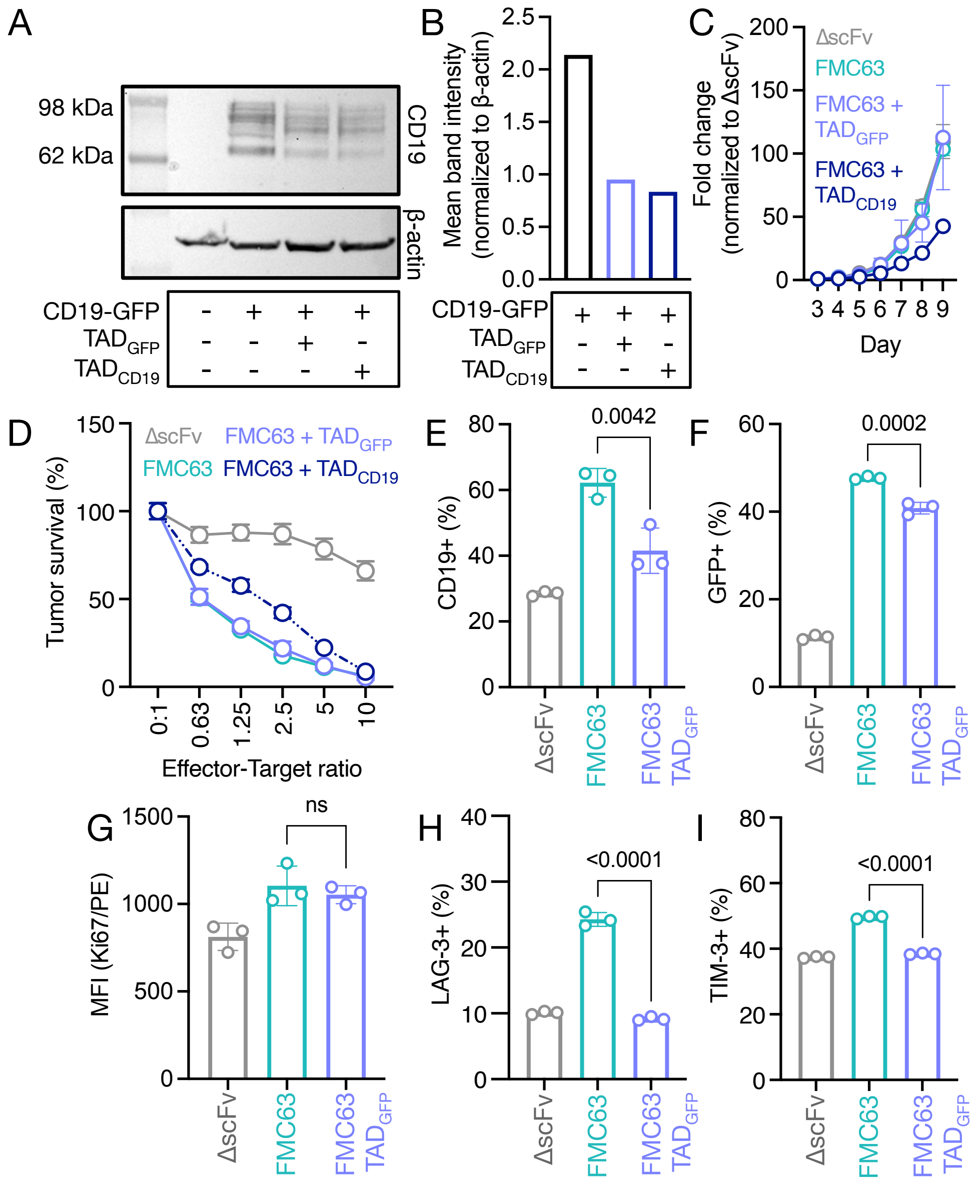


**Supplementary Figure S2: CAR-mediated trogocytosis directly causes CAR T cell dysfunction. (A)** Western blot showing expression of CD19 on 293T cells with or without TAD. Blot is representative of two independent experiments. **(B)** Quantification of western blot in Suppl. Fig. S2A. Values are normalized to β-actin. **(C)** Fold change expansion of FMC63 CAR T cells ± $TAD_{GFP}$/$TAD_{CD19}$ during manufacturing as determined by cell counting, normalized to ΔscFv. Data represents mean ± S.D. from three independent CAR T cell productions using cells from 3 different healthy donors for FMC63 CAR T cells ± $TAD_{GFP}$ and a single production for FMC63 $TAD_{CD19}$ CAR T cells. **(D)** Survival of CD19-GFP-expressing NALM6-luc cells after 16-hour coculture with FMC63 CAR T cells ± $TAD_{GFP}$/$TAD_{CD19}$ or ΔscFv CAR T cells at the indicated effector-target ratios. Tumor survival was measured using a luciferase-based cytotoxicity assay. Data represent mean ± S.D. of three technical replicates and is representative of three independent experiments. **(E)** Percent of CAR T cells positive for CD19 following a 1-hour coculture with CD19-GFP-expressing NALM6 cells as determined by flow cytometry. **(F)** Percent of CAR T cells positive for GFP following a 1-hour coculture with CD19-GFP-expressing NALM6 cells as determined by flow cytometry. **(E/F)** Data is representative of three independent experiments. **(G)** Mean fluorescence intensity of Ki67 in CAR T cells after a 24-hour coculture with CD19-GFP-expressing NALM6 cells as determined by flow cytometry. Data is representative of two independent experiments. **(H)** Percent of CAR T cells positive for LAG-3 after a 24-hour coculture with CD19-GFP-expressing NALM6 cells using full-spectrum flow cytometry. **(I)** Percent of CAR T cells positive for TIM-3 after a 24-hour coculture with CD19-GFP-expressing NALM6 cells using full-spectrum flow cytometry. **(H/I)** Data is representative of three independent experiments. **(E-I)** Data represent mean ± S.D. of 3 technical replicates. Statistical significance was determined by one-way ANOVA.

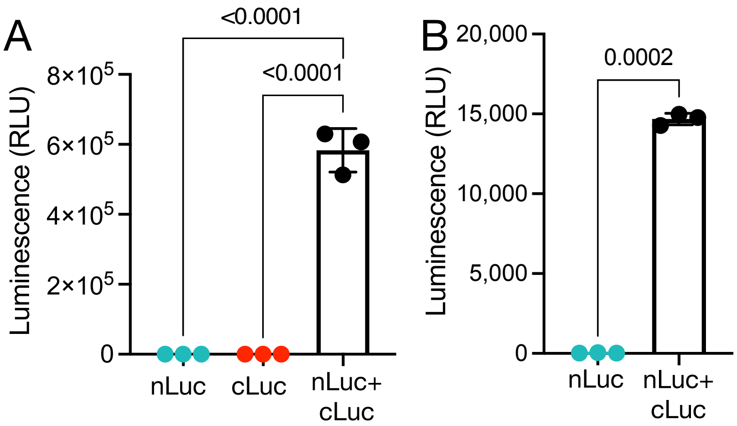


**Supplementary Figure S3: Development of a luciferase complementation assay. (A)** Luminescence of 293T cells 24 hours after transfection with nLuc, cLuc, or nLuc+cLuc containing constructs. Statistical significance was by one-way ANOVA. **(B)** Luminescence of nLuc+ or nLuc+ cLuc+ CAR T cells after manufacturing. Statistical significance was determined by two-tailed Student's *t*-test. **(A/B)** Data represent mean ± S.D. of 3 technical replicates. Data is representative of two independent experiments.

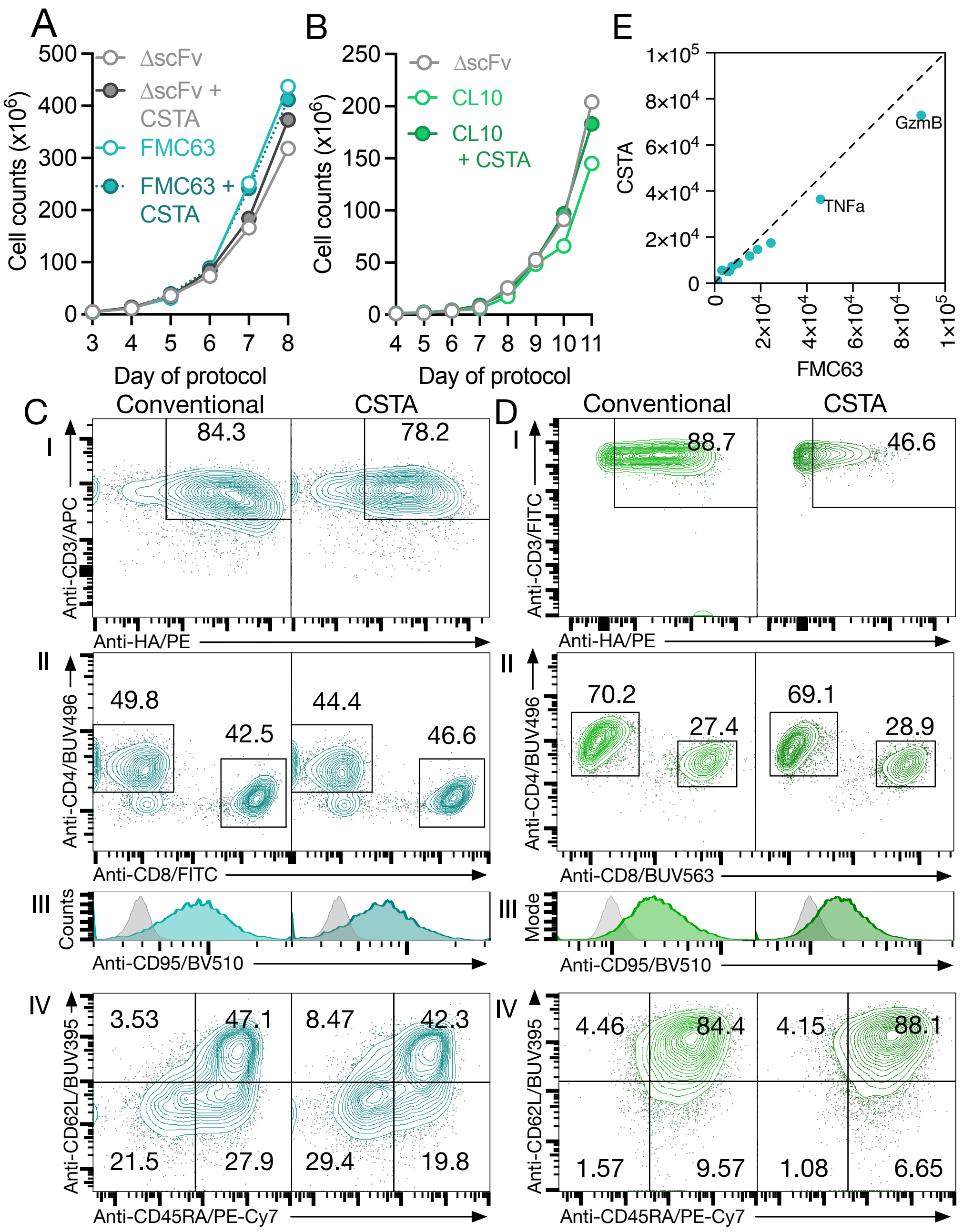


**Supplementary Figure S4: Overexpression of CSTA does not alter CAR T cell expansion, phenotype, or cytotoxic activity. (A)** Expansion of ΔscFv, FMC63 CAR or FMC63 $CAR_{CSTA}$ T cells during manufacturing as determined by cell counting. **(B)** Expansion of ΔscFv, CL10 CAR, or CL10 $CAR_{CSTA}$ CAR T cells during manufacturing as determined by cell counting. **(C-D)** FMC63 (C) or CL10 (D) CAR or $CAR_{CSTA}$ T cells were analyzed by flow cytometry after production for **(I)** CAR expression, **(II)** T cell subsets, and **(III-IV)** T cell phenotype. **(E)** Comparison of secretome between FMC63 CAR and $CAR_{CSTA}$ T cells after 24 hours of activation using anti-CD3/CD28 beads as measured by CodePlex assay. Data represents the average from CAR T cells produced from three healthy donors .

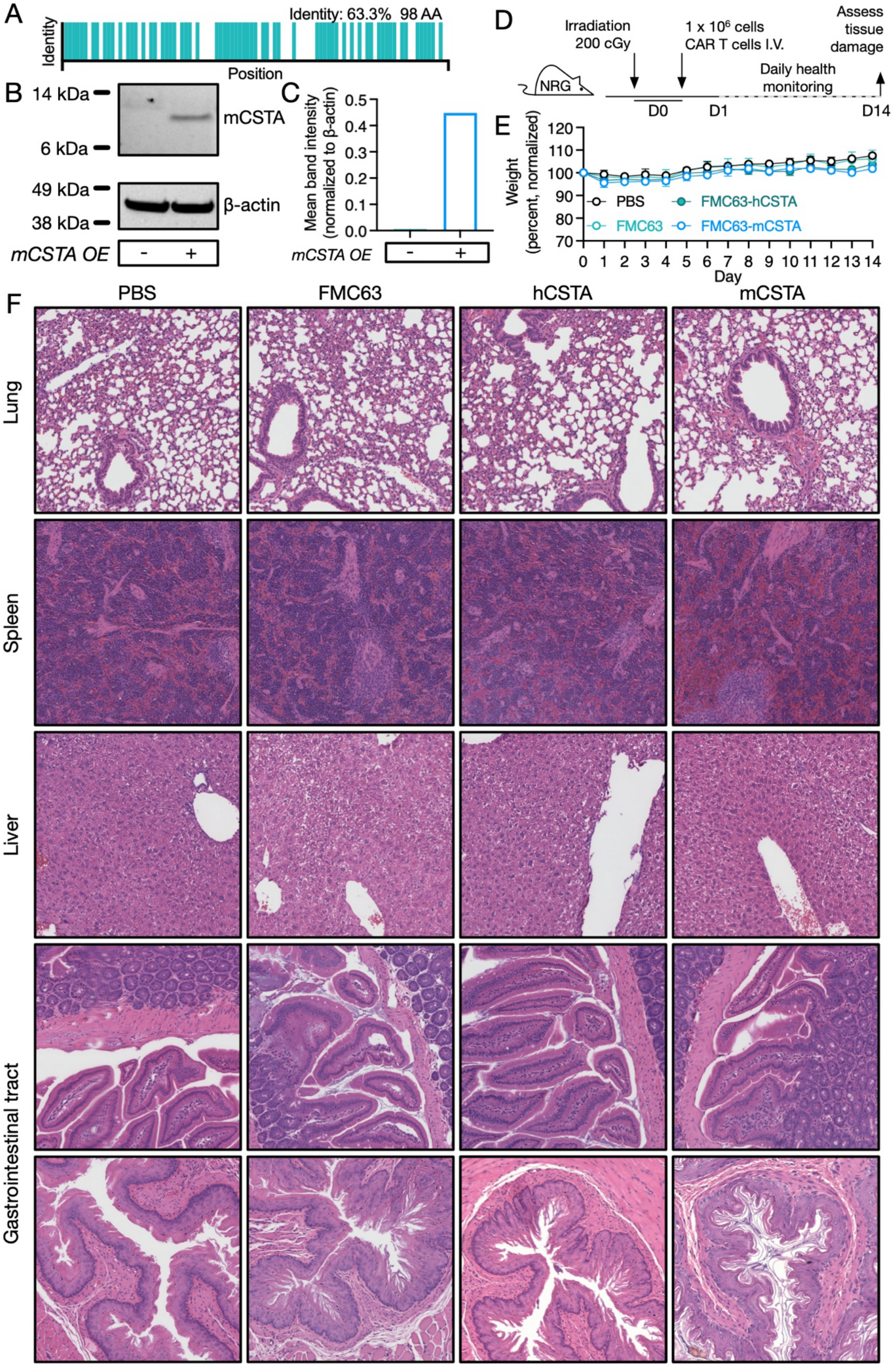
A
Identity: 63.3% 98 AA
Identity
Position
B
14 kDa
6 kDa
mCSTA
49 kDa
38 kDa
β-actin
mCSTA OE
-
+
C
Mean band intensity (normalized to β-actin)
0.5
0.4
0.3
0.2
0.1
0.0
mCSTA OE
-
+
D
Irradiation 200 cGy
1 x 10^6 cells CAR T cells I.V.
Assess tissue damage
NRG
Daily health monitoring
D0
D1
D14
E
Weight (percent, normalized)
120
110
100
90
80
70
PBS
FMC63
FMC63-hCSTA
FMC63-mCSTA
Day
0 1 2 3 4 5 6 7 8 9 10 11 12 13 14
F
PBS
FMC63
hCSTA
mCSTA
Lung
Spleen
Liver
Gastrointestinal tract

62

**Supplementary Figure S5: Overexpression of CSTA does not cause toxicity in mice. (A)** Homology between human cystatin A (hCSTA, Uniprot P01040) and murine cystatin A (mCSTA, Uniprot P56567) as determined by blastp. **(B)** Expression of mCSTA in human FMC63 CAR T cells engineered to overexpress mCSTA as determined by western blot. **(C)** Quantification of western blot in Suppl. Fig. S5B. **(D)** Schema of *in vivo* experiment to assess potential toxicity of hCSTA or mCSTA overexpression. **(E)** Weights of mice treated with ΔscFv, conventional FMC63, or FMC63 CAR T cells engineered to overexpress hCSTA or mCSTA. Data represent mean ± S.D. of 3 mice per group. **(F)** H&E staining of lungs, spleens, livers, and gastrointestinal tracts of mice treated with PBS, conventional FMC63 CAR T cells, or FMC63 CAR T cells overexpressing human CSTA or mouse CSTA. Images are representative of samples from three mice per group.

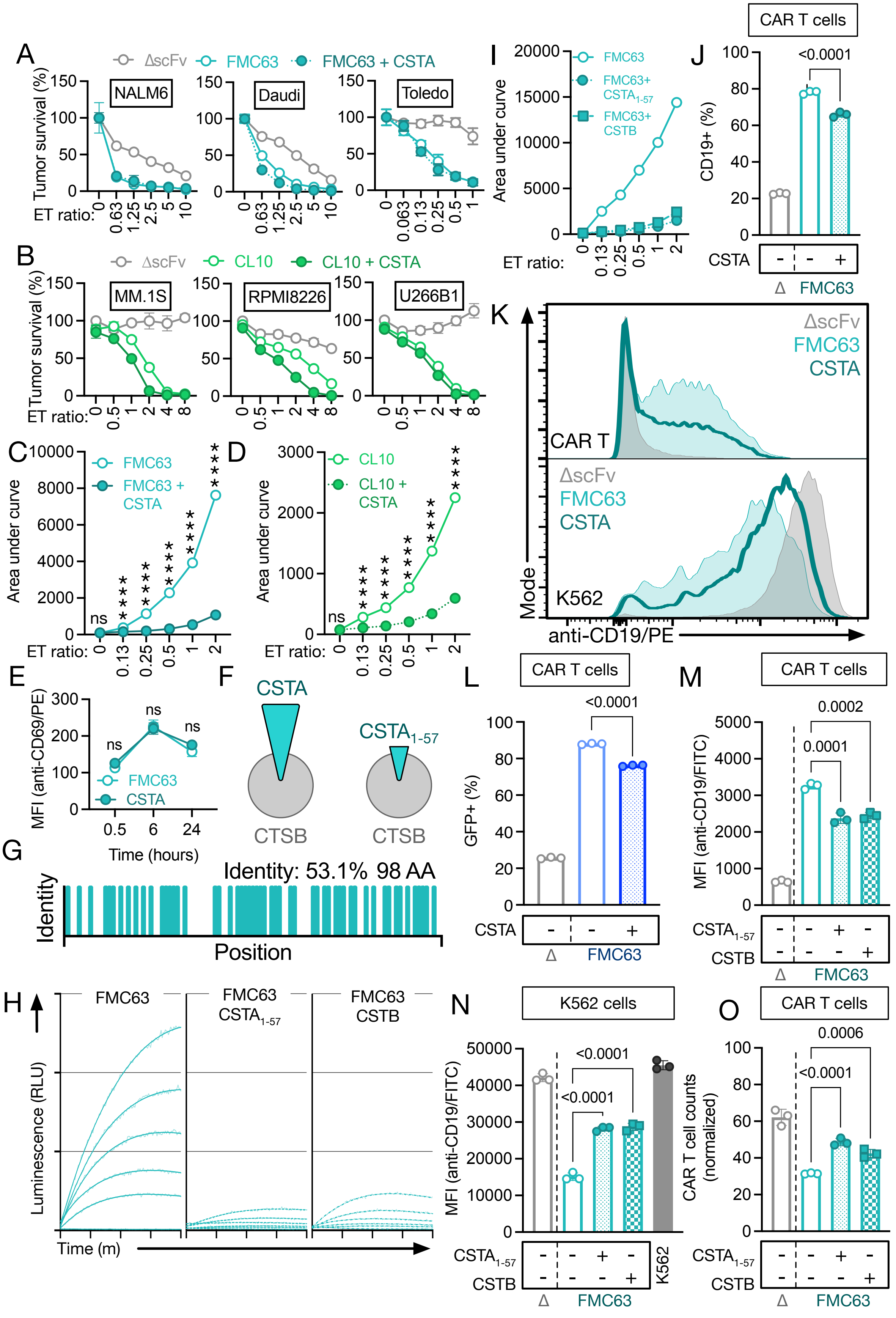
A
ΔscFv FMC63 FMC63 + CSTA
NALM6
Daudi
Toledo
Tumor survival (%)
ET ratio:
B
ΔscFv CL10 CL10 + CSTA
MM.1S
RPMI8226
U266B1
C
FMC63
FMC63 + CSTA
Area under curve
D
CL10
CL10 + CSTA
E
MFI (anti-CD69/PE)
Time (hours)
F
CSTA
$CSTA_{1-57}$
CTSB
G
Identity: 53.1% 98 AA
Identity
Position
H
FMC63
FMC63 $CSTA_{1-57}$
FMC63 CSTB
Luminescence (RLU)
Time (m)
I
FMC63
FMC63+ $CSTA_{1-57}$
FMC63+ CSTB
J
CAR T cells
CD19+ (%)
<0.0001
CSTA
Δ FMC63
K
ΔscFv
FMC63
CSTA
CAR T
K562
Mode
anti-CD19/PE
L
CAR T cells
GFP+ (%)
<0.0001
M
CAR T cells
MFI (anti-CD19/FITC)
0.0002
0.0001
$CSTA_{1-57}$
CSTB
N
K562 cells
<0.0001
K562
O
CAR T cells
CAR T cell counts (normalized)
0.0006

74

**Supplementary Figure S6: Cystatin abundance controls CMT in vitro and in vivo without reducing short-term cytotoxicity. (A)** Survival of NALM6-, Daudi-, and Toledo-luc cells after a 16-hour coculture with FMC63 CAR or $CAR_{CSTA}$ T cells at the indicated effector-target ratios. Data represent mean ± S.D. of three technical replicates. Data is representative of at least three independent experiments using cells produced from three healthy donors. **(B)** Survival of MM.1S-, RPMI8226-, and U266B1-luc cells after 16-hour coculture with CL10 CAR or $CAR_{CSTA}$ T cells at the indicated effector-target ratios. Data represent mean ± S.D. of three technical replicates and is representative of three independent experiments. **(A/B)** Tumor survival was measured using a luciferase-based cytotoxicity assay. **(C)** Area-under-curve quantification of CMT as determined by CompLuc assay in Fig. 5F. **(D)** Area-under-curve quantification of CMT as determined by CompLuc assay in Fig. 5G. **(C/D)** Data represent mean ± S.D. of three technical replicates. Statistical significance was measured using a two-tailed Student's *t*-test. **(E)** Mean fluorescence intensity of CD69 in FMC63 CAR or $CAR_{CSTA}$ T cells using flow cytometry. Statistical significance was measured using a two-tailed Student's *t*-test. Data represent mean ± S.D. of 3 technical replicates. Data is representative of two independent experiments. **(F)** Schema of interaction between CSTA or $CSTA_{1\text{-}57}$ with CTSB. **(G)** Homology between human cystatin A (Uniprot P01040) and cystatin B (Uniprot P04080) as determined by blastp. **(H)** CMT as determined by CompLuc using K562 cells expressing CD19-cLuc after 2 hours. Data represents best fit of three technical replicates and is representative of at least three independent experiments. **(I)** Area-under-curve quantification of luminescence in Suppl. Fig. S6H. Data represent mean ± S.D. of three technical replicates. **(J)** Percent of CAR T cells positive for CD19 following a 30-minute coculture of FMC63 CAR or $CAR_{CSTA}$ T cells and K562 cells expressing CD19-cLuc at a 0.5:1 effector-target ratio as determined by flow cytometry. Data represent mean ± S.D. of 3 technical replicates. Statistical significance was determined using one-way ANOVA. **(K)** Histograms depicting CD19 transfer to CAR T cells following a 30-minute coculture of FMC63 CAR or $CAR_{CSTA}$ T cells and K562 cells expressing CD19-cLuc at a 0.5:1 effector-target ratio as determined by flow cytometry. **(J/K)** Data is representative of at least three independent experiments using cells produced from three healthy donors. **(L)** Percent of CAR T cells positive for GFP following a 1-hour coculture of FMC63 CAR or $CAR_{CSTA}$ T cells and A673 cells expressing CD19-GFP at a 1:1 effector-target ratio as determined by flow cytometry. Data is representative of two independent experiments using cells produced from three healthy donors. **(M)** CD19 transfer to CAR T cells following a 30-minute coculture of FMC63 CAR, $CAR_{CSTA\ 1\text{-}57}$, or $CAR_{CSTB}$ T cells and K562 cells expressing CD19-cLuc at a 0.5:1 effector-target ratio. **(N)** CD19 loss on tumor cells following a 30-minute coculture of FMC63 CAR, $CAR_{CSTA\ 1\text{-}57}$, or $CAR_{CSTB}$ T cells and K562 cells expressing CD19-cLuc at a 0.5:1 effector-target ratio. **(O)** total CAR T cells following a 30-minute coculture of FMC63 CAR, $CAR_{CSTA\ 1\text{-}57}$, or $CAR_{CSTB}$ T cells and K562 cells expressing CD19-cLuc at a 0.5:1 effector-target ratio. CAR T cell numbers are normalized to wells containing only CAR T cells using counting beads. Data represent mean ± S.D. of three technical replicates. **(L-O)** Data represent mean ± S.D. of 3 technical replicates. Data is representative of at least three independent experiments and statistical significance was determined using one-way ANOVA.

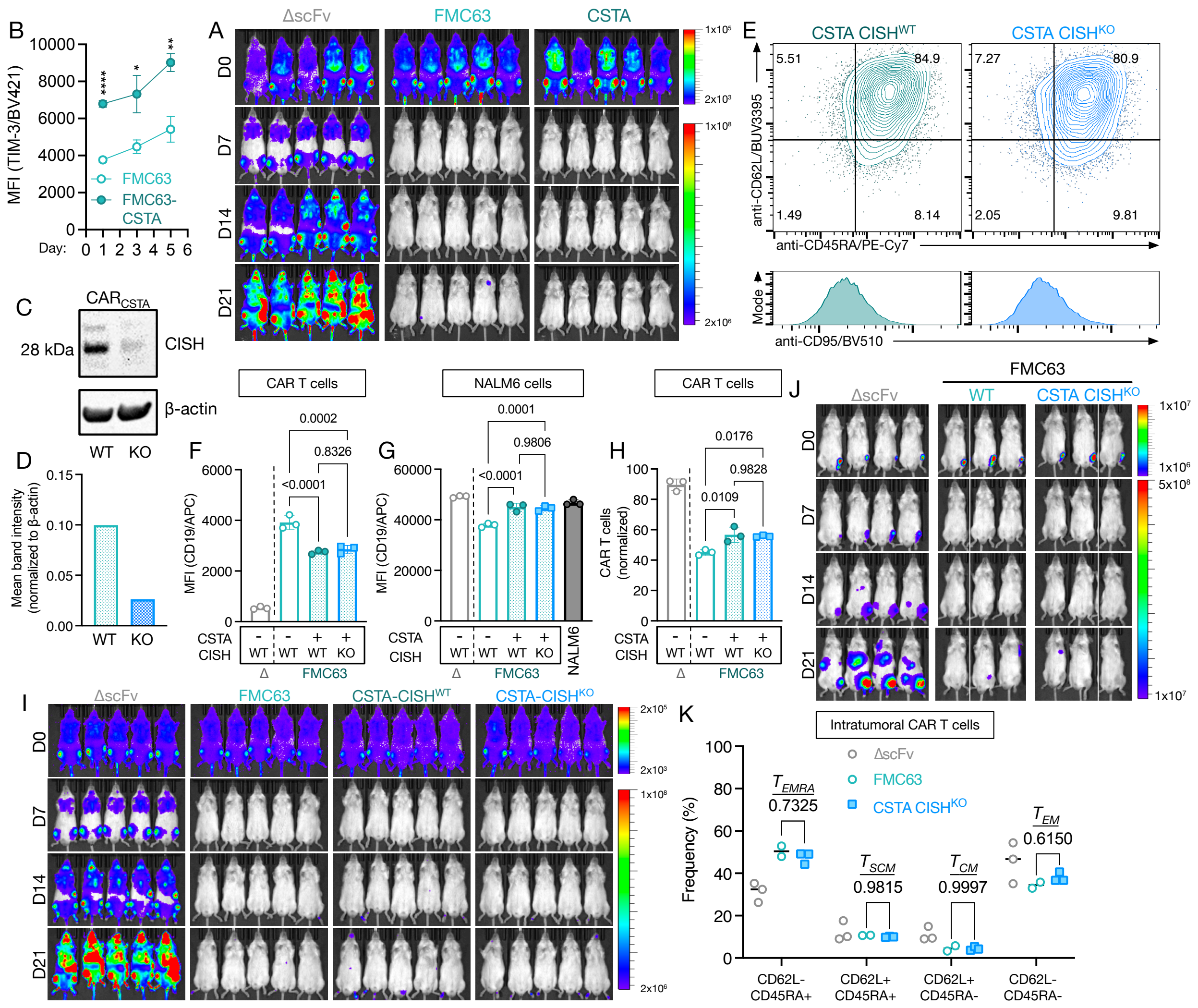


**Supplementary Figure S7: $\text{CAR}_{\text{CSTA}}$ T cells retain benefits of reduced trogocytosis following CISH knockout.** **(A)** Tumor burden in 5 mice per group bearing systemic NALM6 tumors and treated with intravenous ΔscFv, FMC63 CAR, or FMC63 $\text{CAR}_{\text{CSTA}}$ T cells as determined by IVIS. Data is representative of two independent experiments. **(B)** Amounts of TIM-3 in FMC63 CAR or $\text{CAR}_{\text{CSTA}}$ T cells throughout an *in vitro* serial coculture with CD19-GFP-expressing NALM6-luc cells as determined by flow cytometry. Statistical significance was determined using multiple two-tailed Student's *t*-tests. Data represent mean ± S.D. of 3 technical replicates. Data is representative of two independent experiments. **(C)** Western blot showing expression of CISH in $\text{CAR}_{\text{CSTA}}$ $\text{CISH}^{\text{WT}}$ and $\text{CISH}^{\text{KO}}$ T cells. **(D)** Quantification of western blot in Suppl. Fig. S7C. Values are normalized to β-actin. **(E)** Phenotype of FMC63 $\text{CAR}_{\text{CSTA}}$ $\text{CISH}^{\text{WT}}$ or $\text{CISH}^{\text{KO}}$ CAR T cells following production as determined by flow cytometry. **(F)** CD19 transfer to CAR T cells following a 1-hour coculture of FMC63 CAR, CAR$_{CSTA}$ CISH$^{WT}$, or CAR$_{CSTA}$ CISH$^{KO}$ T cells and NALM6 cells expressing CD19-GFP at a 1:1 effector-target ratio as determined by flow cytometry. **(G)** CD19 loss on tumor cells following a 1-hour coculture of FMC63 CAR, CAR$_{CSTA}$ CISH$^{WT}$, or CAR$_{CSTA}$ CISH$^{KO}$ T cells and NALM6 cells expressing CD19-GFP at a 1:1 effector-target ratio as determined by flow cytometry. **(H)** Total CAR T cells following a 1-hour coculture of FMC63 CAR, CAR$_{CSTA}$ CISH$^{WT}$, or CAR$_{CSTA}$ CISH$^{KO}$ T cells and NALM6 cells expressing CD19-GFP at a 1:1 effector-target ratio as determined by flow cytometry. CAR T cell numbers are normalized to wells containing only CAR T cells using counting beads. **(F-H)** Statistical significance was determined using one-way ANOVA. Data represent mean ± S.D. of 3 technical replicates. Data is representative of three independent experiments. **(I)** Tumor burden in 5 mice per group bearing systemic NALM6 tumors and treated with intravenous ΔscFv, FMC63 CAR, or FMC63 CAR$_{CSTA}$ CISH$^{WT}$ or CISH$^{KO}$ T cells as determined by IVIS. **(J)** Tumor burden in 3-4 mice per group bearing intratibial A673 tumors and treated with intravenous ΔscFv, FMC63 CAR, or FMC63 CAR$_{CSTA}$ CISH$^{KO}$ T cells as determined by IVIS. **(K)** Phenotype of intratumoral CAR T cells from mice bearing intratibial A673 tumor cells four days after CAR T cell injection as determined by flow cytometry. Data represent mean ± S.D. from 2-3 animals per group. Statistical significance was determined by two-way ANOVA.

#### SUPPLEMENTARY TABLES

| Target | Flurophore | Clone | Supplier | Target cells |
|---|---|---|---|---|
| BCMA | PE, APC | 19F2 | Biolegend | Multiple |
| CAR detection reagent (FMC63) | PE | n/a | Miltenyi | CAR T cells |
| Cathepsin B | Biotin | n/a | R&D Systems | Western blot |
| Cathepsin L | Biotin | n/a | R&D Systems | Western blot |
| CD3 | APC, FITC, PE, PE-Cy7, PerCP, BV421, BV650 | UCHT1 | Biolegend | Pan-T |
| CD4 | BUV496 | SK3 | BD Biosciences | CD4 T cells |
| CD8 | BUV563, FITC, PE | RPA-T8 | BD Biosciences | CD8 T cells |
| CD19ext | APC, BV650, BV711, PE, FITC, biotin | HIB19 | Biolegend | Multiple |
| CD19int | PE | D4V4B | Cell signaling Technology | Multiple |
| CD20 | APC, BV421, PE-Cy7 | 2H7 | Biolegend | Tumor |
| CD45 | APC | 2D1 | Biolegend | T cells |
| CD45RA | PE/Cy7 | HI100 | Biolegend | T cell subsets |
| CD62L | BUV395, BV421 | SK11 | BD Biosciences | T cell subsets |
| CD69 | PE | FN50 | Biolegend | T cell subsets |
| CD95 | BV510 | DX2 | Biolegend | T cell subsets |
| Cell Trace Far Red | n/a | n/a | Invitrogen | T cells |
| Cytokine-inducible SH2-containing protein | n/a | n/a | Cell signaling Technology | Western blot |
| DAPI | n/a | D1306 | Life Technologies | Live/Dead |
| FolRα | PE | LK26 | Biolegend | Multiple |
| GD2 | APC | 14G2A | Biolegend | Multiple |
| Human Cystatin A | Biotin | n/a | Sino Biological | Western blot |
| Human Cystatin B | n/a | n/a | Sino Biological | Western blot |
| Hemagglutinin | PE, PE-Cy7, APC | 16B12 | Biolegend | CAR T cells |
| HER2t | Alexa Fluor 647 | Hu5 | R&D Systems | CAR T cells |
| Ki67 | PE | Ki-67 | Biolegend | T cell subsets |
| LAG-3 | Alexa Fluor 700 | 11C3C65 | Biolegend | T cell subsets |

| | | | | |
|---|---|---|---|---|
| LINGO1 | Biotin | Opicinumab | Medchemexpress | Multiple |
| Murine Cystatin A | n/a | Polyclonal | Proteintech | Western blot |
| PD-1 | APC-Cy7, PE, PE-Cy7 | EH12.2H7 | Biolegend | T cell subsets |
| TIM-3 | BV421 | 7D3 | BD Biosciences | T cell subsets |
| Zombie Aqua | n/a | n/a | Biolegend | Live/Dead |
| Zombie Red | n/a | n/a | Biolegend | Live/Dead |
| Zombie Violet | n/a | n/a | Biolegend | Live/Dead |
| Zombie NIR | n/a | n/a | Biolegend | Live/Dead |
| Streptavidin | Alexa Fluor 647 | n/a | Jackson Immunoresearch | Multiple |
| β-actin | Unconjugated | 937215 | R&D Systems | Western blot |
| Plasma membrane | Biotracker 555 | n/a | Millipore Sigma | Multiple |

**Supplementary Table S1: Table of monoclonal antibodies and viability dyes used for flow cytometry and western blot analyses.**
